# Nascent dendrite branches initiated by a localized burst of Spire-dependent actin polymerization

**DOI:** 10.1101/2022.06.16.496481

**Authors:** Deirdre Hatton, Claire Marquilly, Caitlin Hanrahan, Tiago Ferreira, Yimiao Ou, Lauren Cinq-Mars, Will Silkworth, Hannah M. Bailey, Margot E. Quinlan, Donald J. van Meyel

**Affiliations:** Centre for Research in Neuroscience, Department of Neurology and Neurosurgery, McGill University, Montreal, Quebec, H3G 1A4, Canada; Research Institute of the McGill University Health Centre, Montreal, Quebec, H3G 1A4 Canada; Integrated Program in Neuroscience, McGill University, Montreal, Quebec, H3A 2B4, Canada; Dept. of Chemistry and Biochemistry, University of California Los Angeles, Los Angeles, CA, 90095, USA; Molecular Biology Institute, University of California Los Angeles, 607 Charles E. Young Drive, Los Angeles, CA 90095, USA

## Abstract

Dendrites form arbors whose size, shape and complexity define how neurons cover their receptive territories. Actin dynamics contribute to growth and remodeling of dendrite arbors. Here we examined how Spire, a conserved actin nucleation factor, promotes formation of new branches in vivo. In live imaging of Drosophila class IV dendritic arborization (c4da) neurons, Spire was observed at new sites of branch initiation, where it assembled new actin polymer in a burst just prior to filopodial outgrowth. For dendrite arborization, Spire required intact structural domains to nucleate actin and target the secretory network, and interacted with Rab11 GTPase, a key regulator of recycling endosomes. Together, these findings support a model in which Spire cooperates with Rab11 to promote new dendrite branches by linking localized actin dynamics with intracellular trafficking of endosomes that deliver lipids and cargoes to fuel protrusive outgrowth of nascent dendrites.

## Introduction

Dendrites have distinct tree-like branching patterns that influence the number, strength, and integration of sensory or synaptic inputs into neurons^1–3^. Understanding how dendrites are shaped during development is essential to gain full insight into the construction of neural circuits that underlie animal survival, behaviors, learning, and memory^4^. A major gap remains in our understanding of the spatial and temporal control of cytoskeletal remodeling responsible for dendrite morphogenesis during development^5^, though it is known to involve regulation of actin dynamics^6^ by specific proteins to nucleate, elongate and remodel actin filaments^7,8^. Furthermore, to cope with their size and complexity, dendrites have specialized arrangements of secretory network organelles for the biogenesis of cargo-carrying vesicles and their transport to the cell surface^9^. Given the importance of actin and the secretory network for dendrite development and function, it is important to clarify how they are coordinated to initiate new dendrite branches in vivo^5,10^.

New dendrite branches arise from actin-rich filopodia that rapidly extend from initiation sites along established branches^10^. Dendrite filopodia are structurally distinct from conventional filopodia, with an unusual network-like cytoskeletal organization characterized by both branched and linear filaments of mixed polarity^11^, suggesting involvement of multiple actin regulatory proteins with distinct activities. Few studies have identified specific actin regulators involved in forming new branches^12–17^, and none have pinpointed their regulation of actin to the location and timing of nascent branch outgrowth. In Drosophila larvae, dendritic arborization (da) neurons provide an excellent model to study the roles of actin regulators in the development of dendrite arbors in vivo^18–20^. In da neurons, the locations of dendrite filopodia initiation sites are pre-figured by patches of actin polymer^12,16,21^, as has been proposed for dendrite filopodia in the mouse brain also^22^. For the large space-filling arbors of class IV da (c4da) neurons, the formation of new dendrites requires the Arp2/3 actin nucleator complex under control of the activator WAVE and the small GTPase Rac1^12^. In c4da neurons, new dendrite formation also involves F-actin severing by Twinstar/Cofilin in the production of “blobs” of pre-assembled actin polymer, which move bidirectionally within dendrites, pause before filopodium outgrowth, and appear to contribute F-actin into the nascent branch^16^. The factors specifying the location and timing of filopodial outgrowth from actin patches and blobs have not been defined.

Here we explored the role of Spire (Spir), a conserved multi-domain actin nucleation factor. We previously showed that knockdown of Spir with RNA interference (RNAi) in cd4da neurons reduced the number of terminal dendrite branches and compromised the ability of these nociceptive neurons to elicit larval escape from noxious stimulus^23^. Spir is known to co-operate with Fmn-family formins, which are also actin nucleators, during Drosophila and mouse oogenesis. Spir has four WASP homology 2 (WH2)-domains for actin nucleation, a KIND domain for interaction with Fmn-family formins^24–27^, a modified FYVE (mFYVE) domain to target vesicles of the secretory network, and a SPIR-box with potential to interact with Rab GTPases^24,28–31^, which play roles in vesicle sorting and transport^32,33^. Previous studies showed functional links between Spir and Rab11 in oocytes, in cultured cells, and in vitro^28,31,34,35^, and Rab11 has been shown to be required for correct dendrite arborization of c4da neurons^36–38^. Here we show that Spir stimulates a burst of localized actin polymerization just prior to filopodial outgrowth of new dendrite branches and that Spir cooperates with Rab11 for dendrite arborization. Our data supports a model in which Spir and Rab11 link localized actin assembly with the secretory network to deliver lipids and cargoes to nascent dendrites.

## Results

### Spir is required in c4da neurons for localized F-actin synthesis just prior to outgrowth of new dendrite branches

Development of Drosophila c4da dendrites begins in embryogenesis and continues in larval stages, with arbors growing in the larval body wall between muscles and epithelial cells^18^. We produced a *GAL4* line inserted in the *spir* locus (*spir^GAL^*^4^) and used it to drive the reporter *UAS-mCD8::GFP*. *spir^GAL4^* was expressed in c4da neurons (marked by ppk- CD4::tdTom) and other sensory neurons in the larval body wall (Fig. 1A), consistent with our previous report^23^. To examine Spir protein expression within c4da neurons, we developed new anti-Spir sera for immunohistochemistry (IHC) (Fig. S1A), and we also introduced an in-frame epitope-tag (smGFP-HA) into the *spir* locus (Fig. S1C, D). Both approaches showed Spir to be expressed in c4da neurons in third instar (L3) larvae.

**Figure 1:**
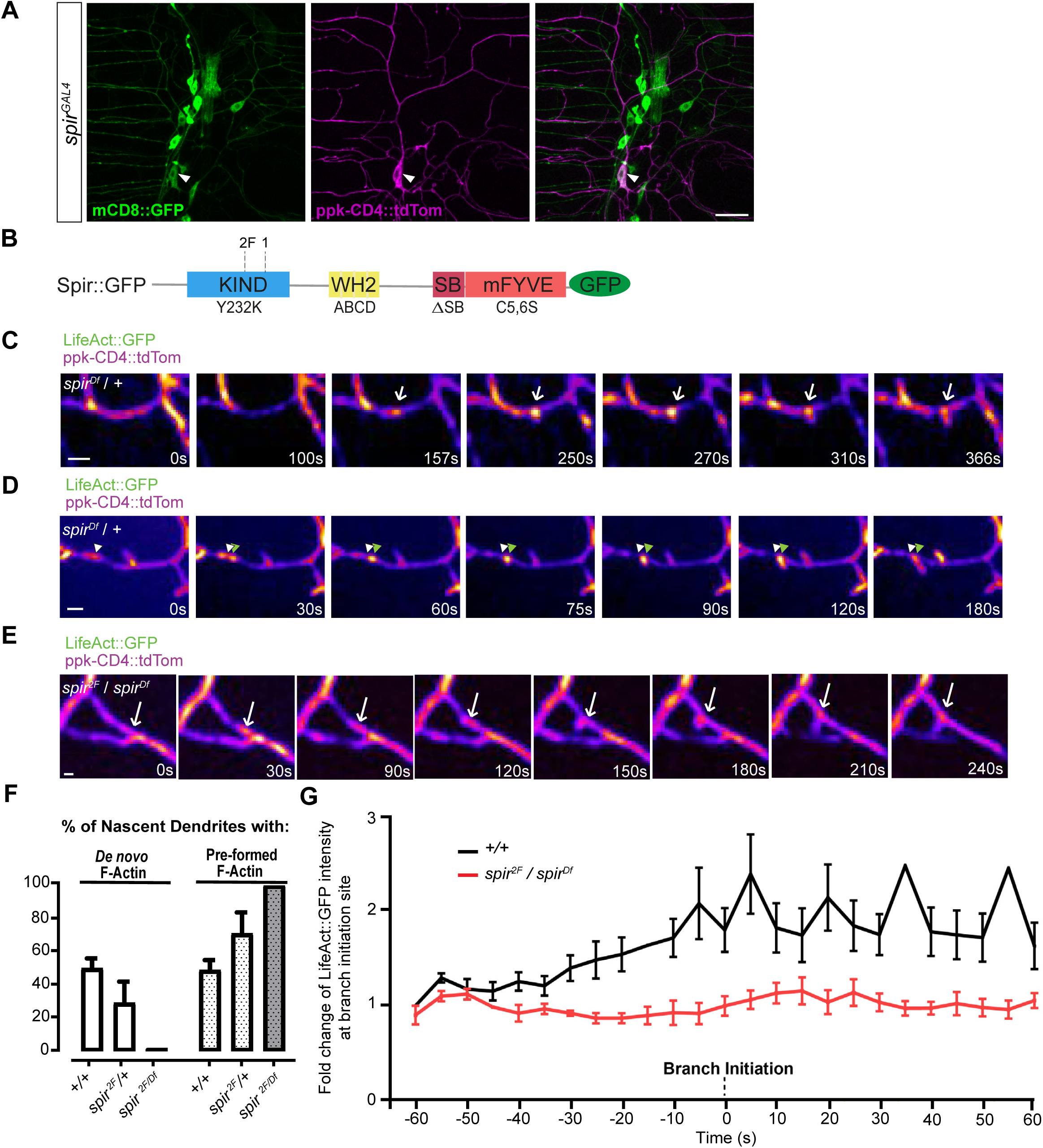
Spir is required for localized F-actin synthesis just prior to new branch outgrowth. (A) *spir^GAL4^*-driven mCD8::GFP in c4da neurons (ppk-CD4::tdTom) including ddaC (arrowhead), and other sensory neurons in an L3 larva. Scale bar=25μm. (B) Diagram showing Spir protein domains (KIND, WH2, SB, and mFYVE), the locations *spir^2F^* and *spir^1^* mutations, and the names of the disruptive mutations (Y232K, ABCD, ΔSB, and C5,6S) introduced into Spir::GFP. (C, D) Filmstrips (Supplemental Videos 1 and 2) of new dendrite branch formation in a c4da neuron (ppk-CD4::tdTom) from a control (*spir^Df^*/+) L2 larva. LifeAct::GFP (driven by *ppk-GAL4*) reveals actin polymer within dendrites. Arrow (in C) points to a punctum of F-actin appearing *de novo*, and arrowheads (in D) point to a pre-existing punctum that moved from its original position (white arrowhead) to the site of branch initiation (green arrowhead). In all instances, LifeAct::GFP signal intensified just prior to branch initiation. Scale bar = 2μm. (E) Filmstrip (Supplemental Video 3) showing new dendrite branch formation in a *spir^2F^*/*spir^Df^* larva at L2, where LifeAct::GFP moved to the initiation site but did not intensify prior to outgrowth (arrow). Scale bar = 2μm. (F) Proportion of newly formed terminal branches in L2 larvae that had either *de novo* F-actin or pre-formed F-actin prior to branch initiation in c4da neurons. Graph shows mean ± SEM, comparing controls (+/+, n=50 new branches in movies from 11 neurons) with *spir* heterozygotes (*spir^2F^*/+, n=18 new branches, 8 movies) and *spir* mutants (*spir^2F^*/*spir^Df^*, n=8 new branches, 11 movies). (G) Fold change of F-actin intensity (LifeAct:GFP) at branch initiation sites, from 60s before branch formation to 60s afterward, comparing mean ± SEM of controls (black) and *spir* mutants (red).

To reveal actin dynamics leading up to the protrusion of dendrite filopodia, we acquired time-lapse movies in second instar (L2) larvae, capturing F-actin (labeled with LifeAct::GFP) within c4da neurons (labeled with ppk-CD4::tdTom). To study how loss of *spir* affects actin accumulation and new branch initiation in c4da neurons, we used the mutant alleles *spir^1^* and *spir^2F^*, which prematurely truncate Spir within the KIND domain (Fig. 1B)^39^. We examined each allele in combination with a deficiency for *spir* (*spir^Df^*), which has a distinct genetic background. All heteroallelic combinations of these alleles of *spir* are viable, and so with anti- Spir sera we were able to confirm that *spir* mutants lack Spir expression in larval sensory neurons (Fig. S1B), and adult brain lysates (Fig. S1E).

We captured movies of developing c4da dendrites in L2 larvae and mapped the appearances of new branches in control animals heterozygous for *spir^Df^* (Fig. 1C, D) or *spir^2F^/+* (Fig. 1F), or in wild-type (+/+, Fig. 1F). In all of these controls LifeAct::GFP was seen in stationary puncta that appeared prior to branch initiation; in wild-type (+/+) approximately half of these puncta appeared *de novo* (Fig. 1C, F) and the other half were pre-formed actin blobs that moved to the site (Fig. 1D), consistent with a previous report^16^. In *spir^2F^/spir^Df^*mutants, however, new branches arose only from pre-formed actin blobs that moved to the initiation site and not from stationary actin puncta produced *de novo* (Fig. 1E, F).

In controls where branches initiated from either blobs or stationary puncta, LifeAct::GFP signal at these sites became more intense just prior to branch initiation and then extended into the protruding nascent branchlet (Fig. 1C, D). We quantified this change of intensity in wild-type (+/+) controls, where LifeAct::GFP increased roughly two-fold on average in the 60s prior to outgrowth and remained at this intensity for ≥ 60s (Fig. 1G). In contrast, in *spir^2F^/spir^Df^* mutants the LifeAct::GFP signal in blobs did not increase prior to branch initiation (Fig. 1E, G), indicating that Spir is required for this localized assembly of F-actin just prior to the outgrowth of new dendrite branches.

### Localization of Spir at nascent branches

In c4da dendrites, Spir was observed in puncta in dendrite branches near the cell body, but strong labelling from epithelial cells and muscles prevented their detection and quantification in more distal branches (Fig. S1A, C, D). To overcome this, we expressed a *UAS-Spir::GFP* transgene (Fig. 1B) in c4da neurons with *ppk-GAL4,* and characterized the distribution of Spir::GFP (Fig. S1F). In L3 larvae, GFP-labeled puncta were seen in 100% of shafts, 28% of branch points, and 6% of branch tips (Fig. S1G). Consistent with this, most of these GFP-labeled puncta were found along shafts (57%) or at branch points (36%) (Fig. S1G). We next used live imaging to determine if Spir::GFP localized to sites of new branch initiation in L2 larvae. Movies (∼1 frame/2 seconds (sec) for >5 minutes (min)) showed a distribution of Spir::GFP (Fig. 2A, B) similar to that of fixed larvae at L3 (Fig. S1F). Some puncta were stationary while others were steadily motile or moved in a saltatory manner with occasional pauses (Fig. S1H). Motile puncta were often smaller and rounder, while stationary puncta were often larger and irregular in shape. Importantly, every nascent branch in controls (+/+, L2) had a punctum stationed precisely at (68%) or near (< 2 µm, 32%) the initiation site prior to outgrowth (Fig. 2C). These puncta arrived from elsewhere (30%, Fig. 2A, D) or were already at that site in the first frame (70%, Fig. 2B, D), and time until branch outgrowth varied (Fig. 2D). Spir::GFP intensity did not typically increase prior to outgrowth but, in some instances (12%), a motile punctum combined with one already stationed at the future initiation site. Spir::GFP at branch initiation sites was stably maintained throughout the process of filopodial outgrowth, stabilization, or retraction, and invaded the filopodia as they grew (Fig. 2B).

**Figure 2:**
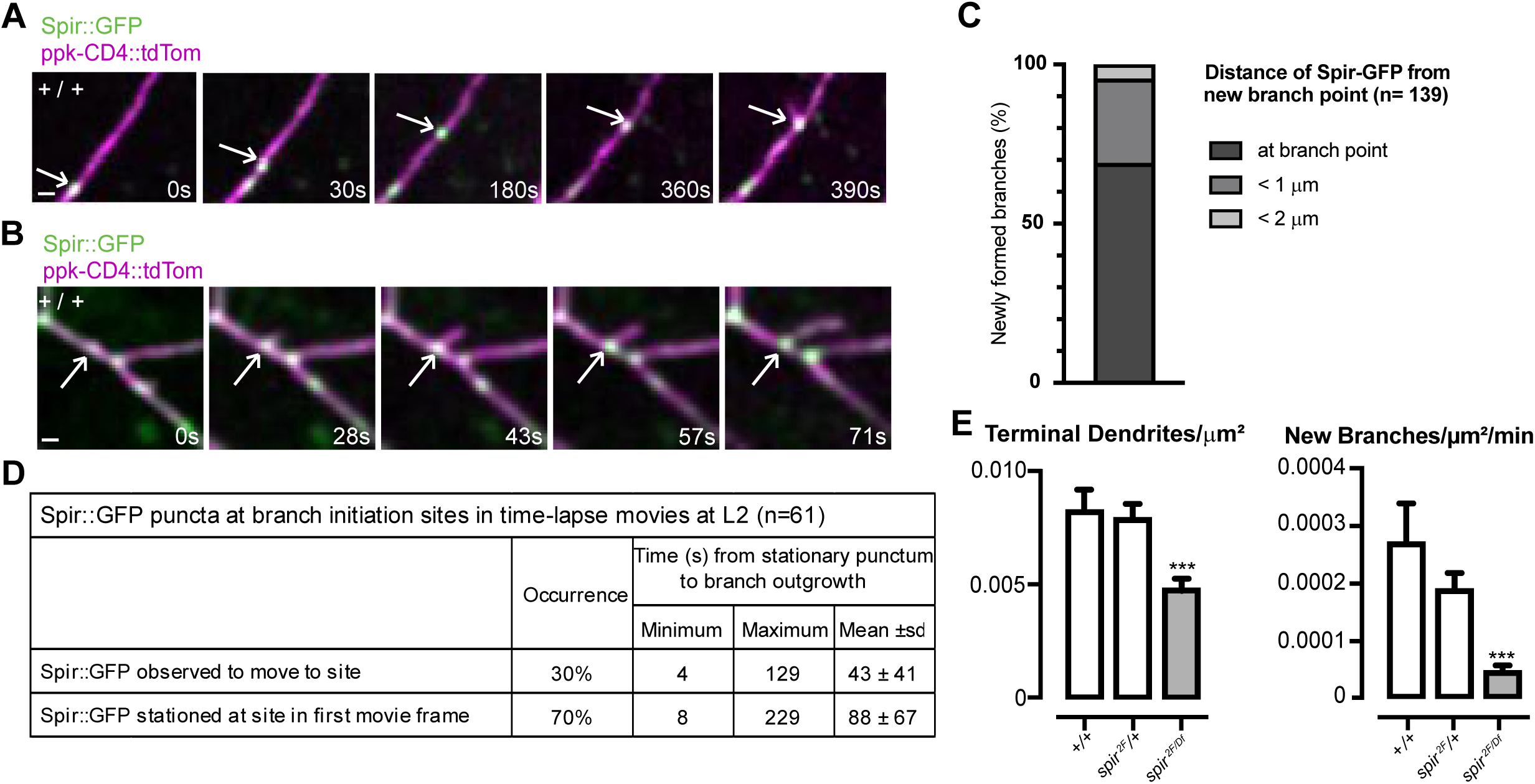
Spir is at initiation sites for new dendrite branches, and is required for their formation. (A, B) Filmstrips (Supplemental Video 5 and Supplemental Video 6) of control (+/+) L2 larva, where Spir::GFP (arrows) was observed to move to the initiation site of a new dendrite branch (A) or was already stationed there at the beginning of the movie (B). Scale bars = 2μm. (C, D) For newly formed dendrite branches, the distance of the closest Spir::GFP puncta from the site of branch initiation was measured (C), and the time from observation of a stationary Spir::GFP punctum at the site to filopodial outgrowth was recorded (D). (E, F) Dendrite parameters from time lapse movies of control (+/+, n=104 new branches, 10 movies), heterozygotes (*spir^2F^*/+, n=91 new branches, 10 movies) or *spir^2F^*/*spir^Df^*mutants (n=42 new branches, 15 movies). (E) Terminal branches/µm^2^ in first movie frame (ANOVA F(2,32)=12.30, p<0.0001). (F) Number of new branches/µm^2^/min in each movie (ANOVA F(2,32) =9.276, p=0.0007).

### Spir is required for new branch formation

At the outset of our timelapse movies, the number of terminal dendrites/µm^2^ was reduced in *spir^2F^/spir^Df^* mutants compared to controls and, importantly, so was the number of new branches/µm^2^ formed per min during each movie (Fig. 2E). This was also observed in movies of c4da neurons expressing LifeAct::GFP instead of Spir::GFP (Fig. S2A, B). For new branches, growth occurred by saltatory extensions in both controls and *spir^2F^/spir^Df^* mutants, but there were no differences between them in mean speed of outgrowth, proportion of time spent paused between extensions, or maximum length of protrusion (Fig. S2C-E). For pre-existing branches, the proportion that were static increased in *spir^2F^/spir^Df^* mutants (Fig. S2F- I), whether LifeAct::GFP was present or not. Together these results confirm that Spir promotes the formation of new branches and contributes to dynamics of pre-existing ones.

Consistent with this, we examined the consequences of Spir function for fully formed arbors of c4da neurons at L3 and found that *spir* mutants (*spir^1^/spir^Df^* or *spir^2F^/spir^Df^*) had fewer terminal branches than controls (*spir^2F^*/+) (Fig. 3A, C), and thus shorter total arbor length (Fig. 3D). Sholl analysis also showed reduced peaks of maximum branch density (critical value) (Fig. 3B, F). Dendritic field was reduced in some c4da neurons of s*pir^2F^/spir^Df^* mutants (Fig. 3E), and so we repeated the analysis after removing a subset (4/15) of cells with unusually small dendritic fields. As expected, we found that dendritic field was no longer significantly different from controls (Fig. S3A) but the branching deficits were still strongly evident for the number of terminal dendrites, arbor length and critical value (Fig. S3A-D).

**Figure 3:**
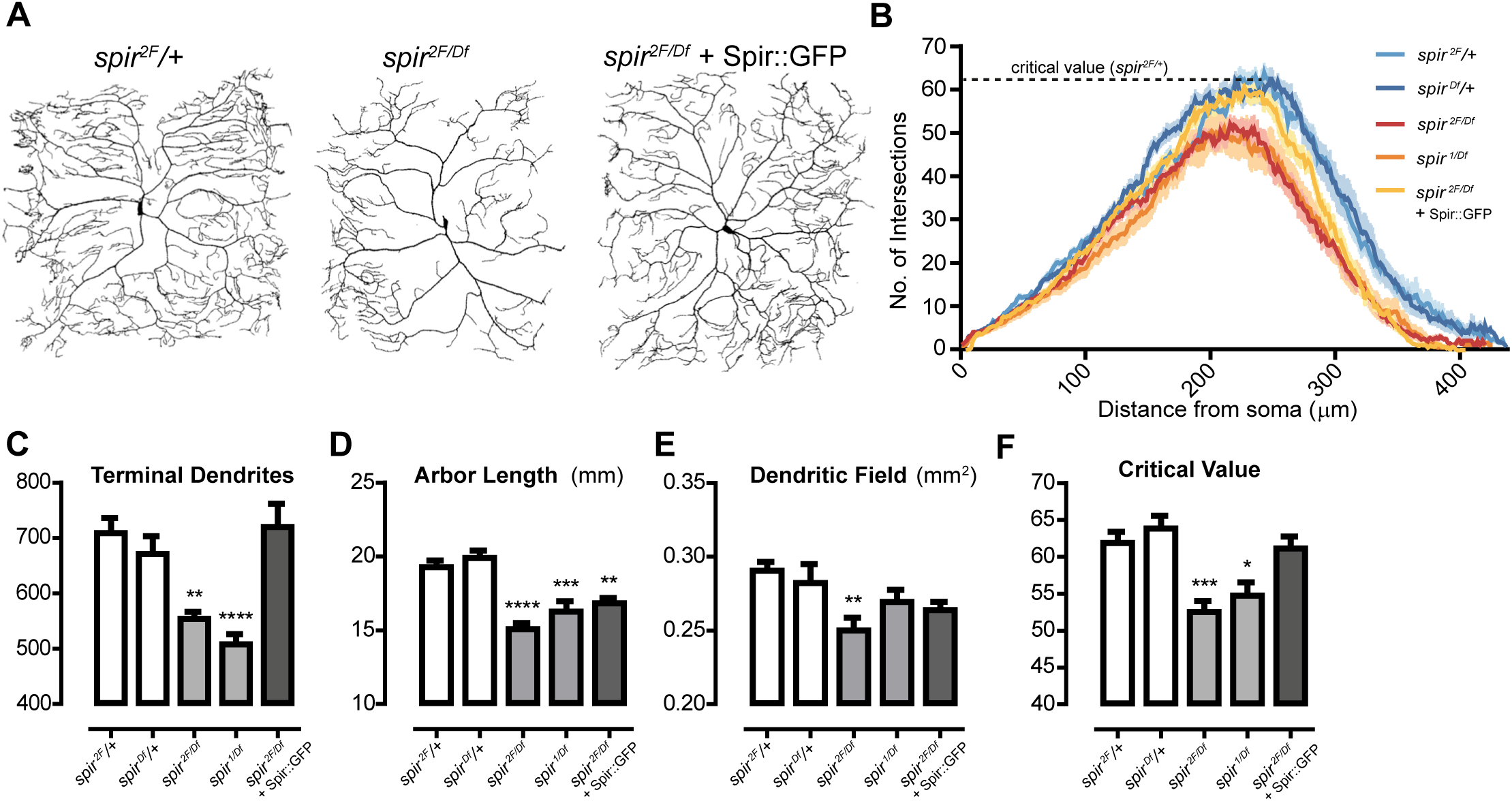
Spir is required for dendrite arborization of c4da neurons. (A) Dendrite arbors in a *spir* heterozygote control (left), a *spir* mutant (middle), and a *spir* mutant rescued with Spir::GFP (right). (B) Sholl profiles with dotted line indicating critical value for controls (*spir^2F/+^*). (C-F) Dendrite parameters per neuron in L3 larvae (n=15 for each genotype). Asterisks indicate significant changes compared to *spir^2F/+^* heterozygous controls. (C) Total number of terminal dendrites per neuron (ANOVA (F(4, 70) = 11.30, p < 0.0001). (D) Total length of dendrite arbor (mm) (ANOVA F(4, 70) = 15.12, p < 0.0001). (E) Dendritic field (mm^2^) (ANOVA (F(4, 70) = 3.352, p = 0.0144). (F) Sholl critical value (ANOVA (F(4, 70) = 8.936, p < 0.0001).

Importantly, the selective expression of *UAS-spir::GFP* in c4da neurons with *ppk- GAL4* (Fig. 3A) fully rescued number of terminal branches in *spir^2F^/spir^Df^* mutants (Fig. 3C, Fig. S3B), and critical value (Fig. 3B, F, Fig. S3D), demonstrating that Spir acts within c4da neurons to control these features. It did not fully rescue total arbor length (Fig. 3D, Fig S3C), which is not likely to be due to inadequate expression since IHC (Fig. S1F) and western blots (Fig S1E) showed Spir::GFP at higher levels than endogenous Spir. Perhaps additional cell types contribute to total arbor length, such as epithelial cells in the larval body wall where Spir is expressed (Fig. S1A-D). We conclude that Spir acts within c4da neurons to establish the correct number of terminal branches, likely by contributing to filopodial outgrowth necessary for nascent branches.

### Each domain of Spir is required for dendrite arborization

We sought to determine the importance of the four identified protein domains of Spir by testing if disruptive mutations in Spir::GFP (Fig. 1B) could rescue the correct numbers of terminal branches in c4da neurons of *spir* mutants. The Y232K mutation is a missense substitution in the KIND domain known to abolish the interaction between Spir and the Fmn-family formin Cappuccino (Capu) in oocytes^25,40^. The ABCD mutation is a series of 4 triple-alanine substitutions in the WH2 domains that greatly diminish the ability of Spir to polymerize F-actin *in vitro*^24^. The ΔSB mutation is a 20 aa deletion of the entire Spir-box domain which has sequence similarity to the Rab effector protein Rabphilin3a suggesting it interacts with Rab GTPases^28^. Finally, the C5,6S mutation involves substitution of serine residues for 2 key cysteines in the so-called modified FYVE (mFYVE) domain. FYVE domains interact in lipid membranes with Phosphatidylinositol 3-phosphate (PtdIns3P), a prominent component of the membranes in the endosomal system^41^. The mFYVE domain in Spir binds less specifically to negatively charged lipids, including PtdIns3P, and is required for Spir’s localization to intracellular membranes^29^.

When expressed in c4da neurons (with *ppk-GAL4)* of *spir^2F^/spir^Df^*mutants, none of these mutations rescued the number of terminal dendrites per neuron (Fig. 4A) or critical value (Fig. 4B, Fig. S4A). This was not due to insufficient supply because none were expressed at lower levels than intact Spir::GFP in western blots of adult head lysates (Fig. 4C, D and Fig. S4B). The Y232K, ABCD, and C5,6S mutations did not obviously affect the punctate distribution of Spir::GFP in static images of c4da arbors in L3 larvae (Fig. S4C), but the ΔSB mutation caused it to accumulate near the cell soma and form a more continuous pattern (Fig. 4E). Though this might be explained in part by unusually high expression levels (Fig. 4C, D and Fig. S4B), these results suggest that the SB domain is required for proper localization of Spir within c4da arbors. None of the mutants appear to have dominant-negative activity, since expression in a wild-type (+/+) background caused no significant differences in number of terminal dendrites or total arbor length compared to Spir::GFP (Fig. S5A-E), though the ΔSB mutation trended toward reductions of both parameters. Based on the failure of these four mutations to rescue *spir^2F^/spir^Df^* mutants, we conclude that each domain of Spir contributes to its control of dendrite branching.

**Figure 4:**
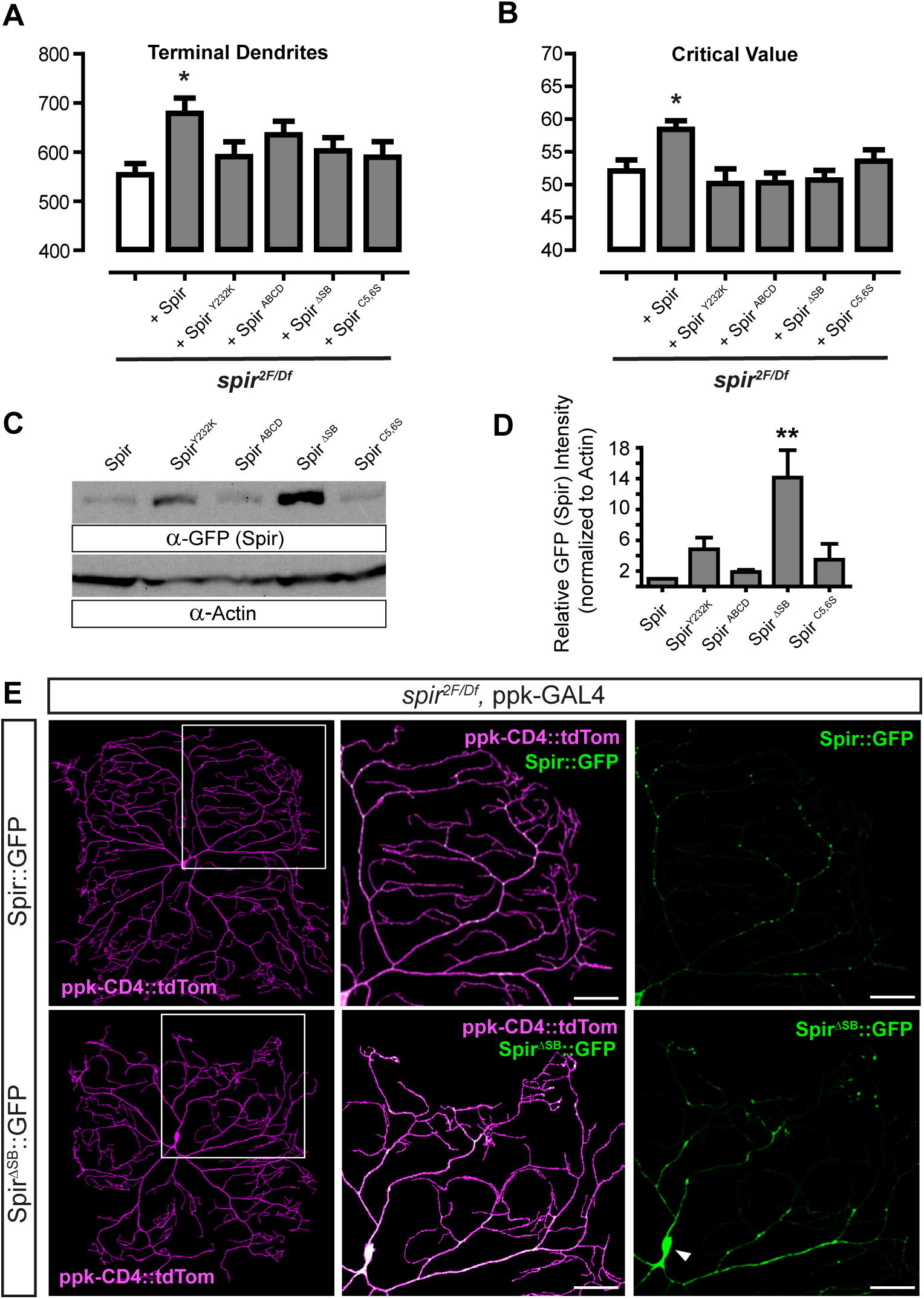
Structure-function analysis to identify Spir domains required for dendrite arborization and Spir localization. (A-B) Dendrite parameters (L3 larvae) showing rescue of *spir^2F^*/*spir^Df^* mutants with Spir::GFP, but failure to rescue by domain-specific mutations (n=15 for each genotype). (A) Total number of terminal dendrites per neuron (ANOVA (F(5, 83) = 2.269, p = 0.0550). (B) Sholl critical value (ANOVA (F(5, 84) = 3.552, p = 0.058). (C) Anti-GFP and anti-Actin western blot analysis from adult head lysates, where transgenes were expressed in all neurons with nSyb-GAL4. Expected size of Spir::GFP is 142 kDa, and endogenous Actin is 42 kDa. (D) Relative band intensities on western blots (n=3) of Spir::GFP constructs (anti-GFP) normalised to anti-Actin. Spir^ΔSB^::GFP is expressed at significantly higher levels than Spir::GFP (Friedman test (Q (4) = 10.40, p = 0.0053), with Dunn’s post-hoc tests). (E) *ppk-GAL4*-driven Spir::GFP or Spir^ΔSB^::GFP in a *spir* mutant background. White boxes in left panels indicate areas magnified in panels to their right. Scale bar=50μm. Spir::GFP is punctate in dendrites, but Spir^ΔSB^::GFP accumulates in a continuous pattern in dendrites proximal to the cell soma (white arrowhead).

### Spir and Rab11 cooperate for dendrite arborization

The pattern and motility of Spir::GFP puncta within dendrites of c4da neurons was reminiscent of elements of the secretory network, where Rab GTPases including Rab11 are key regulators of vesicle and cargo trafficking^32^. Rab11 is associated with the trans-Golgi network, post-Golgi, and recycling endosomes^42^. Rab11 has been shown to be required for correct c4da dendrite arborization^36–38^, a finding we confirmed with RNAi knockdown of *Rab11* mRNA in c4da neurons (Fig. 5A) which reduced the numbers of terminal dendrites in L3 larvae (Fig. 5B), total arbor length (Fig. 5C), dendritic field (Fig. 5D), and Sholl critical value (Fig. 5E, F) compared to controls (+/+). In contrast, overexpression of Rab11::GFP in c4da neurons was not sufficient to affect the number of terminal dendrites, arbor length, and Sholl critical value at L3 (Fig. S6A, B, D), though the dendritic field was smaller (Fig. S6C).

**Figure 5.**
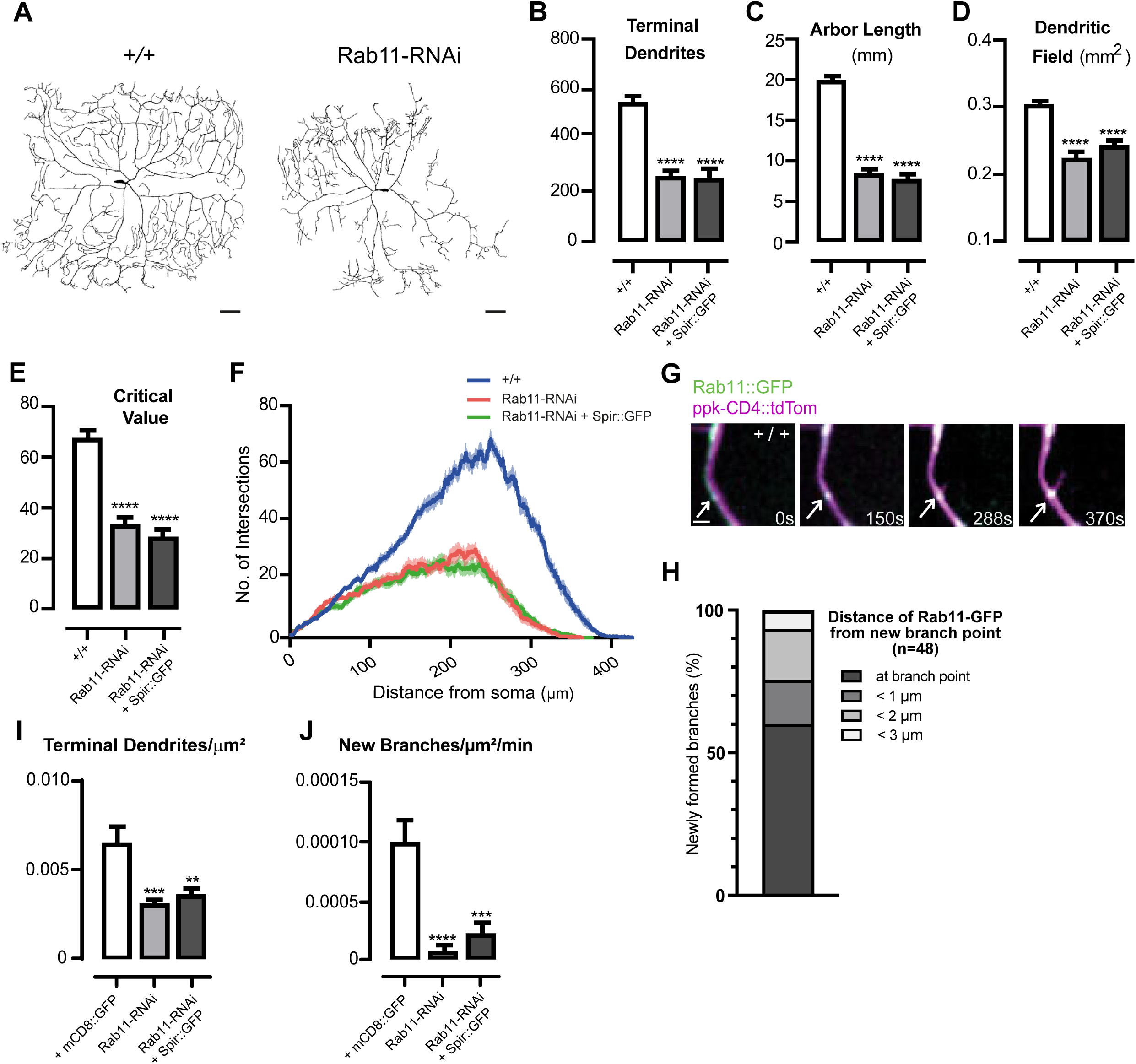
Rab11 is required in c4da neurons for dendrite arborization and new branch formation. (A) Dendrite arbors of c4da neurons in control (+/+) (left) and *ppk-Gal4*-driven Rab11-RNAi (right) L3 larvae. Scale bar=50μm. (C-G) Dendrite arbor parameters in control (*+*/+, n=10), *ppk-Gal4*-driven Rab11-RNAi (n=10), and *ppk-Gal4*-driven Rab11-RNAi and Spir::GFP (n=12). Asterisks indicate significant changes compared to +/+ controls. (B) Total number of terminal dendrites (ANOVA (F(2,29) = 41.06 p<0.0001). (C) Total length of dendrite arbor (mm) (ANOVA (F(2,29) = 163.6, p<0.0001). (D) Dendritic field (mm^2^) (ANOVA (F(2, 29) = 34.00, p<0.0001). (E) Sholl critical value (ANOVA (F(2, 29) = 79.84, p<0.0001) (F) Sholl profiles. (G) Filmstrip (Supplemental Video 7) of control (+/+) L2 larva, where Rab11::GFP (arrows) was observed at the initiation site of a new dendrite branch. Scale bar = 2µm. (H) Distance of the closest Rab11::GFP puncta from the site of branch initiation (n=48) in control (+/+) L2 larvae. (I, J) Dendrite parameters from time lapse movies of control (+mCD8::GFP, n=72 new branches, 10 movies), *ppk-Gal4*-driven Rab11-RNAi (n=6 new branches, 11 movies) or *ppk-Gal4*-driven Rab11-RNAi and Spir::GFP (n=18 new branches, 9 movies). (I) Terminal branches/µm^2^ in first movie frame (ANOVA F(2,27)=12.41, p=0.0002). (J) Number of new branches/µm^2^/min in each movie (ANOVA F(2,27) =14.29, p<0.0001).

Whether Rab11 is involved in the initiation of new dendrite branches has not been determined. To address this, we mapped the appearance of new branches in movies (∼1 frame/2 sec for >5 min) of developing c4da dendrites in L2 larvae (Fig. 5G). We began by examining the distribution of Rab11::GFP, the expression of which had no effect on the dynamics of pre-existing branches (Fig. S6E-H). In controls (+/+, L2) we observed Rab11::GFP at (60%) or near (< 3 µm, 40%) the initiation site of each new dendrite branch (Fig. 5H). Some Rab11::GFP puncta were motile within arbors, but for every nascent branch we observed (n=48), Rab11::GFP was already stationed at the initiation site from the beginning of every movie and so we could not determine how long it had been there prior to outgrowth. Depletion of Rab11 by RNAi knockdown reduced the total number of terminal branches/µm^2^ at L2 (Fig. 5I) and, importantly, severely reduced the number of new branches/µm^2^/min in each movie (Fig. 5J), indicating that Rab11 is indeed required for the initiation of new dendrite branches. As in *spir* mutants (Fig. S2F-I), pre-existing branches were also affected by Rab11 knockdown (Fig. S6E- H): the proportion of branches that both extended and retracted was reduced (Fig. S6G), while the proportion of static branches increased (Fig. S6H). Therefore, loss of Rab11 results in fewer new branches and reduced dynamics of existing ones.

Since Rab11, like Spir, is required for proper dendrite arborization, is localized to sites of new branches, and is required for their initiation, we wondered how Rab11 and Spir might be related in regulating these processes. We first asked whether expressing excess Spir::GFP could restore dendrite arborization and new branch formation to neurons lacking Rab11. We found that it did not (Fig. 5B-F, I, J). In the converse experiment, Rab11::YFP overexpression did not restore dendrite arborization to *spir* mutants (Fig. S6I-M). Together these findings suggest that Spir and Rab11 do not lie in a simple linear pathway controlling dendrite development, and so we wondered whether Spir and Rab11 might interact and thereby cooperate with one another for dendrite arborization instead.

Genetic interaction experiments that examine trans-heterozygotes for mutant alleles of *spir* and *Rab11* can provide evidence that they cooperate in a common molecular pathway. When compared to control heterozygotes for either mutation alone (Fig. 6A) the c4da neurons of trans-heterozygotes showed reduced numbers of terminal branches (Fig. 6B) and arbor length (Fig. 6C), while dendritic field was largely unaffected (Fig. 6D).

**Figure 6.**
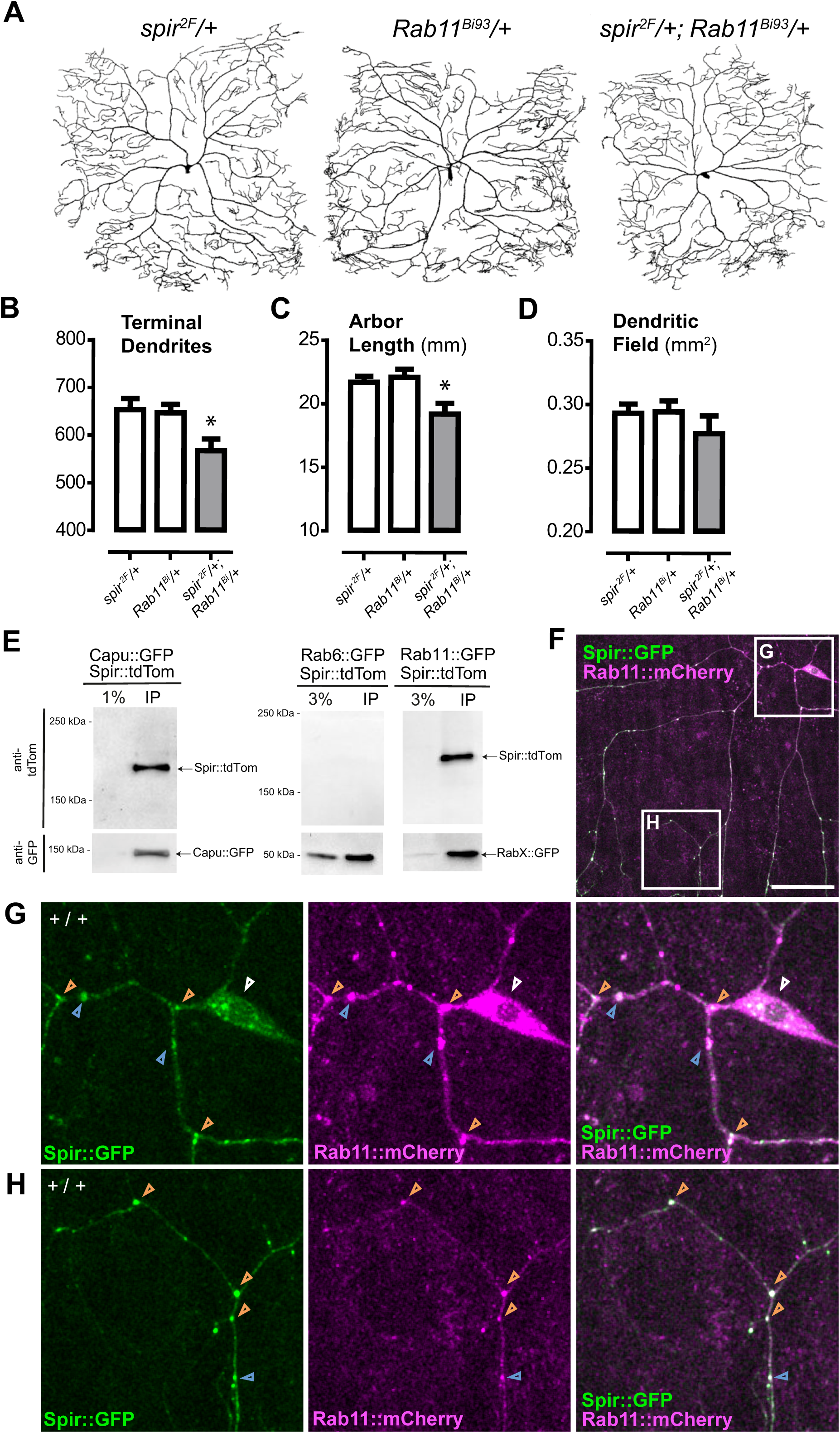
Spire interacts with Rab11 in c4da neurons. (A) Dendrite arbors of c4da neurons in *spir^2F^/+* heterozygotes (left), *Rab11^Bi93^* heterozygotes (middle), and trans-heterozygotes for both (right). (B-D) Dendrite arbor parameters in *spir* heterozygotes (*spir^2F^*/+, n=15), *Rab11* heterozygotes (*Rab11^Bi93^*/+, n=17), and trans- heterozygotes (*spir^2F^*/+; *Rab11^Bi93^*/+, n=15). Asterisks indicate significant changes compared to *spir^2F^/+* heterozygous controls (*P<0.05). (B) Total number of terminal dendrites (ANOVA (F(2, 44) = 4.745 p = 0.0136). (C) Total length of dendrite arbor (mm) (ANOVA (F(2, 44) = 5.443, p =0.0077). (D) Dendritic field (mm^2^) (ANOVA (F2, 44) = 0.8512, p=0.4338). (E) Spir::tdTom co-immunoprecipitates with either Capu::GFP or Rab11::GFP from adult head lysates, but not with Rab6::GFP. In SDS-PAGE gels, GFP-Trap® immunoprecipitates (IP) were run alongside 1% or 3% of initial lysate (as indicated), and replicate immunoblots were probed with antibodies to detect GFP or tdTom. (F-H) *ppk-GAL4*-driven Spir::GFP (green) and Rab11::mCherry (magenta). (F) Boxes indicate regions of a c4da neuron selected for magnified view of the cell body (white arrowhead) and proximal dendrites (G) or distal dendrites (H). There are many instances of co-localization of Spir::GFP and Rab11::mCherry at branch points (e.g., orange arrowheads) and dendrite shafts (e.g., blue arrowheads). Scale bar in F=50μm.

We next tested whether Spir and Rab11 can physically associate with one another *in vivo*, by co-expressing Spir-tdTom (in all neurons, with *nSyb-GAL4*) with either Rab11::GFP, or Capu::GFP (a positive control), or Rab6::GFP (a negative control). From lysates of adult heads, Spir::tdTom co-immunoprecipitated with Rab11::GFP and with Capu::GFP, but not with Rab6::GFP (Fig. 6E). This suggests that Spir could selectively associate with Rab11- containing protein complexes in neurons. Consistent with this, we found that upon simultaneous expression of Rab11::mCherry and Spir::GFP in c4da neurons, they were usually co-localized in the cell body, at dendrite branch points and along dendrite shafts (Fig. 6F-H).

Since Spir and Rab11 co-localized extensively in dendrites, we wondered whether the positioning of one might be dependent on the other. Upon Rab11 knockdown, the punctate distribution of Spir::GFP was unaffected at L3 (Fig. S7A, B) and, in live imaging at L2, Spir::GFP was still positioned at (56%) or near (< 2 µm, 44%) each new branch initiation site (n=18, Fig. S7C, D). Therefore, Rab11 did not appear to be required for Spir localization at branch initiation sites, and so we wondered if Spir might be required for Rab11 localization instead. For technical reasons, we were unable to examine either Rab11::GFP or Rab11::YFP in *spir* mutants, and so we studied the marker Sec15::GFP instead. Sec15 is a Rab11 binding protein ^43–45^ and a component of the exocyst, a Rab effector complex of eight proteins involved in the delivery of material to the plasma membrane ^46,47^. Like Rab11::GFP, Sec15::GFP showed punctate, motile expression in dendrite arbors in live imaging at L2, where Sec15::GFP was positioned at (80%) or near (< 1 µm, 20%) each initiation site (n=21 new branches in 8 movies, Fig. S7E,F). Sec15::GFP was observed at initiation sites in rare instances where nascent branches were captured in *spir* mutants (n=2, from 11 movies, Fig. S7G), suggesting Spir is not required for positioning Sec15::GFP.

We have found that Spir and Rab11 are both required for the synthesis of new dendrite branches and, furthermore, that they genetically interact in vivo, physically interact in neurons and co-localize in dendrites. Together this data provides strong evidence that Spir and Rab11 functionally interact with one another, either directly or indirectly, in a common molecular pathway for dendrite arborization in c4da neurons.

## Discussion

Spir was identified previously as being regulated by Longitudinals Lacking (Lola), a BTB/POZ transcription factor involved in dendrite development^23^ and motor axon guidance^48^. However, the mechanisms of Spir function in neuronal development have remained unclear. Our data presented here indicate that Spir stimulates localized assembly of actin in the seconds prior to outgrowth of new dendrite branches, and that Spir function is linked to the GTPase Rab11 in the formation of new branches and the dynamic motility of existing ones. Unlike Spir::GFP which arrived from elsewhere in 30% of new branch initiation events, Rab11::GFP was already at the site from the beginning of every movie, suggesting that, at least in some cases Spir is recruited to a pre-existing site marked by the presence of Rab11. Rab11 has been shown to regulate movement of recycling endosomes toward the plasma membrane^49^, and the trafficking of receptors and trans-membrane proteins necessary for neurite outgrowth and remodeling ^50,51^. Sec15 is a cofactor for Rab11 and a subunit of the exocyst complex, which mediates the tethering of secretory vesicles to the plasma membrane prior to fusion and has been shown to be required for c4da dendrite growth, maintenance, and regeneration following injury^52,53^. We therefore propose that Spir cooperates with Rab11 to couple actin dynamics with the delivery of lipids and cargoes through the secretory network to the exocyst complex, fueling protrusive outgrowth of nascent dendrites. Consistent with this, Spir function in dendrite arborization required intact domains for actin assembly (KIND, WH2) and for targeting Spir to the secretory network (mFYVE). The Spir-box has potential to mediate interactions with Rab GTPases, and we found that it was essential to control the levels and localization of Spir in dendrites. However, the functional and physical interaction between Spir and Rab11 did not involve their interdependent localization, raising the possibility that other factors are involved in the recruitment of Spir to nascent branch initiation sites within arbors.

In control animals, newly formed branches were pre-figured either by the appearance of actin polymer *de novo* or by the arrival of pre-formed actin blobs, in roughly equal proportions. In *spir* mutants, there were fewer new branches, and none stemming from initiation sites with *de novo* actin polymer, indicating that Spir is critical for new dendrite formation from such sites. It is unclear whether stationary actin patches and motile actin blobs are features of convergent or parallel processes to initiate dendrite branches^54^. Although Spir was present and active at initiation sites stemming from pre-formed actin blobs, it was dispensable for branch initiation from at least some of them. From this we infer redundancy with other molecular mechanisms for branch initiation, perhaps involving actin synthesized by the Arp2/3 complex which has been shown to localize transiently to branch initiation sites in c4da neurons^12^.

Spir::GFP overexpression did not affect dendrite arborization (compare Fig. 3 and Fig. S5), supporting the idea that its expression at nascent branch sites reflects that of endogenous Spir. Since Spir::GFP overexpression was not sufficient to induce ectopic branching, and the interval between stationing of Spir at a branching site and filopodial outgrowth was variable, our findings suggest that Spir activity is tightly regulated and may require additional partners to engage the switch to branch initiation. The KIND domain mutation Y232K is known to abolish the interaction between Spir and the formin Capu, and there is some evidence that Spir and Capu interact for the spike-like protrusions in dendrites of class 3 da neurons^17^. However, in unpublished data we built a *capu^GAL4^* line that did not label c4da neurons, and RNAi lines targeting Capu did not reduce terminal branch number when expressed in c4da neurons. Of note, the KIND domain also binds intramolecularly to the mFYVE domain^29^, and so perhaps the Y232K and C5,6S mutations in these domains disrupt this interaction and thereby impact Spir function in dendrites. Alternatively, Spir might cooperate with another formin in c4da neurons such as Formin3, which is required for c4da dendrite arborization^55^. Further research will be required to identify a formin partner for Spir in c4da neurons, or to determine if Spir could nucleate actin on its own in neurons, as has been shown *in vitro*^24^.

Interestingly, the Formin3 ortholog INF2 has been reported to interact with mouse Spir- 1 in mammalian cells^56^. Murine orthologs of Spir (Spir-1 and Spir-2) are expressed in the developing CNS and adult brain^57,58^. Decreased Spir-2 expression is associated with epilepsy in humans and mice^59^, but it is not known if either Spir-1 or Spir-2 are involved in dendrite arborization in mammals. A distinct actin nucleator known as Cordon-bleu has been demonstrated to promote dendrite branch initiation in cultured hippocampal neurons and slice preparations of cerebellar Purkinje neurons^15^.

Actin filaments generated by Spir proteins and formins have been shown to provide tracks for myosin-V motors in microtubule-independent transport of Rab11-labeled endosomes to the plasma membrane of oocytes^34,60^ and in dispersal of Rab27a-labeled melanosomes in melanocytes^61^. Interestingly, mammalian orthologs of myosin Vb have been shown to bind directly to the orthologs of both Spir and Rab11^35^, which together activate myosin Vb and stimulate its motility^31^. In dendrites, Spir could contribute to an actin pool for myosin-V-mediated delivery of Rab11 vesicles to nascent branches, supplying lipids and proteins required for their protrusive growth. This actin pool could allow for the subsequent invasion of microtubules into these nascent branches, perhaps mediated by myosin-6^62^ or augmin^63^, and the transition to microtubule-dependent long-distance transport of cargo-carrying vesicles. While further experiments are required to fully elucidate the mechanisms of dendrite branching, our findings show that Spir-mediated actin dynamics link branch initiation with protrusive outgrowth in vivo.

## Acknowledgements

This work was supported by the Canadian Institutes of Health Research (FRN137034 to DJvM), National Institutes of Health (R01GM096133 to MEQ), a Ruth L. Kirschstein National Research Service Award (GM007185 to HMB), and studentships from the RI-MUHC and McGill (to DH). The authors thank Greg Emery, Peter McPherson and Martine Girard for advice and sharing reagents, our colleagues Emilie Peco, Eunjoo Cho, Adela Ralbovska, Keith Murai, Sabrina Chierzi and Todd Farmer for helpful suggestions. For technical support, the authors thank Dr. Min Fu and Shi-bo Feng of the RI-MUHC Molecular Imaging Platform, and Tim McLean. Stocks obtained from the Bloomington Drosophila Stock Center (NIH P40OD018537) were used in this study.

## Author Contributions

D.J.v.M and M.E.Q. conceived the project and designed experiments with input from all other authors. D.H., C.M., C.H., T.F., Y.O., L.C-M., W.S., and H.M.B. performed experiments and analyzed all data. D.J.v.M wrote the manuscript with input from all other authors.

## Materials and Methods

### *Drosophila* stocks and genetics

Fly stocks were obtained from the Bloomington Drosophila Stock Centre (*spir^1^*, *spir^2F^*, *spir^DfExel60^*^46^ (referred to as *spir^Df^*), Mi05646, Mi05737, *UAS-capu::GFP*, *Rab11^Bi93^, w^11^*^18^, *UAS-mCD8::GFP*, *UAS-Rab11::GFP*, *UAS-Rab11::YFP, UAS-Rab6::GFP*, *UAS-Rab11-dsRNA* (referred to as *UAS-Rab11-RNAi*), *UAS-LifeAct-GFP*. *ppk-GAL4* and *ppkCD4::tdTom* and *ppkCD4::GFP* were gifts from Dr. Yuh-Nung Jan. *UAS- Rab11::mCherry* and *UAS-sec15-GFP were* a gift from Dr. Gregory Emery. *nSyb-GAL4* line was created by Dr Julie Simpson, for which we used 2nd chromosome insertions generated by Dr Stefan Thor. To create the lines *UAS-spir::GFP*, *UAS-spir::tdTom*, *UAS-spir^Y232K^::GFP*, *UAS-spir^ABCD^::GFP*, *UAS-spir^ΔSB^::GFP*, *UAS-spir^C5,6S^::GFP*, DNA constructs encoding full- length Spir or each of the 4 mutated versions ^24,28,40^ were designed to add C-terminal GFP or tdTomato (tdTom) fluorophores. These constructs were subcloned into the vector pMUH, then microinjected by standard procedures (Bestgene, Inc) to target the VK00027 landing site in the *Drosophila* genome with φc31-mediated integration^64^. *spir^GAL4^*was generated by inserting a so-called Trojan-GAL4 into the *spir* locus^65,66^. pT-GEM(0) a gift from Benjamin White (Addgene plasmid # 62891) was microinjected (Bestgene, Inc) into MiMIC line, Mi05646^65,66^. The same plasmid was injected into MiMIC line Mi05737 to generate *capu^GAL4^*. CRISPR/Cas9- mediated homologous recombination was used to introduce spaghetti-monster GFP-HA (smGFP-HA)^67^ into the *spir* locus. To generate Spir-smGFP-HA flies, we used an approach combining dual guide RNA (gRNA) sequences^68^ and short homology arms^69^. The CRISPR Optimal Target finder (http://targetfinder.flycrispr.neuro.brown.edu/index.php) was used to select gRNA sequences for the C-terminus of Spir (5’- GTCGGCCCTGGATCTGACGCCCGTC-3’) and just after the 3’UTR (5’-GTCGGCAAACTAAAGAACAAGATTC-3’), and these oligonucleotides were cloned into pCFD3-dU6:3gRNA (Addgene plasmid #49410)^70^. Homology sequences on 200 nucleotides (PAM removed) were cloned into a self-linearizable Puc573 vector for homologous recombination^69^. The repair template included smGFP-HA, the endogenous 3’UTR of Spir, and the fluorescent eye reporter 3xP3-dsRed, all flanked by two PiggyBac recombinase sites. Plasmids expressing the gRNAs (100ng/µL) and donor template (250ng/µL) were mixed and micro-injected into nos-Cas95 embryos^71^ (BDSC 78782). Progeny were screened via fluorescence for the 3xP3-dsRed reporter, and fly stocks were established. All injections and initial screening were completed by BestGene (Chino Hills, CA). Proper insertion of smGFP was confirmed via genomic PCR (primers 5’-GGGGATTCAACCTGTTCTCCT-3’ and 5’- TGTGCAAGTGCGTTCTGAAG-3’), and expression confirmed by western blot.

### Antibodies

To produce anti-Spir antisera, DNA encoding Spir amino acids 565-692 were cloned into pGEX-6P-1 to produce an IPTG-inducible C-terminal GST fusion proteins in BL21 cells. The purified GST fusion protein was injected into two rabbits and the serum was extracted according to approved protocols at the Comparative Medicine and Animal Resources Centre (McGill University) and standards established by the Canadian Council on Animal Care. Crude serum (500 µl) was then pre-absorbed by overnight incubation with 500 µl L3 larvae (*spir^1^/spir^Df^*) that had been fixed previously in 4% PFA. Other primary antibodies for IHC were Alexa-647-conjugated goat anti-HRP (1:800, Jackson ImmunoResearch #123605021) and rabbit anti-HA (1:300, Cell Signaling Technology #3724), while the secondary antibody for anti-Spir was Alexa-647-conjugated goat anti-rabbit (1:1000, Invitrogen, #A21244), and for anti-HA IHC it was Alexa-405-conjugated goat anti-rabbit (1:1000, Invitrogen, #A48254). For western blotting, membranes were incubated with mouse anti-GFP (Clontech #632380), rabbit anti-DsRed (Takara #632496, for tdTom detection), mouse anti-Actin (Sigma #A4700) or rabbit anti-Spire (directed at amino acids 565-692). For detection, HRP-conjugated secondary antibodies (anti-mouse and anti-rabbit (both 1:3000, BioRad) were used and revealed with chemiluminescence (Amersham ECL Western Blotting Detection Reagents).

### Immunohistochemistry (IHC)

L3 larvae of either sex were dissected in Sorensen phosphate buffer (pH 7.4) and fixed in 4% paraformaldehyde (in Sorensen’s buffer) for 10 minutes (min) at room temperature. Washes (3X over 20 min) were done with Sorensen’s buffer containing 0.2% Triton-X-100 (Sigma). Specimens were then blocked with Sorensen’s buffer containing 5% normal goat serum (NGS), incubated at 4°C with primary antibodies overnight, washed as above, then incubated with secondary antibodies for 2 hours at room temperature. Anti-Spir antiserum for immunohistochemistry was directed at amino acids 1-91. Primary and secondary antibodies were diluted in Sorensen’s buffer containing 5% NGS. After a final wash as above, specimens were then mounted with SlowFade Diamond Antifade Mountant (Invitrogen) for imaging.

### Co-immunoprecipitation (co-IP) and western blotting

Lysates for co-IPs and western blots were made from adult heads aged 1-5 days after eclosion. For western blots, heads were mechanically crushed using a motorized pestle directly in 2X Laemmli buffer. For co-IPs, 100 heads were cut and collected on dry ice, then mechanically crushed using a motorized pestle in 100ul of lysis buffer containing 50mM Tris pH, 150mM KCl, 1mM MgCl2, 1mM EGTA, Roche protease inhibitor cocktail and 10% glycerol. Lysates were treated with 10% deoxycholic acid and 10% Triton X-100, then incubated overnight on a Nutator with anti-GFP bound agarose beads (Chromotek GFP-Trap® #gta). Lysates and beads were separated in a centrifuge, washed with a glycerol free buffer, and the complexes on the beads were dissociated directly in Laemmli buffer, then separated using standard SDS-PAGE and wet-transfer procedures.

### Imaging

All imaging and quantification was of ddaC, the dorsal-most c4da neuron, in abdominal segments of the larval body wall. For static images, L3 larvae were dissected and prepared for IHC, or they were squished directly on the slide in 100% glycerol and prepared for imaging of native fluorescence (GFP, tdTom, mCherry, mRFP) within 10 minutes. Static images were taken with an Olympus Fluoview FV-1000/BX63 confocal microscope using either a 20X or 60X oil-immersion objective lens. All time-lapse imaging was done on L2 larvae, where eggs were initially collected for 4h on agar plates containing grape juice and yeast. After 24h, hatched larvae of selected genotypes were transferred on a fresh fly food plate to start timing. L2 larvae (65h ± 4h AEL) were mounted in a drop of halocarbon oil in a polydimethylsiloxane (PDMS) microfluid device, and then imaged by live confocal microscopy (Zeiss, Spinning Disk, Inverted Axio Observer Z1) with 63x objective lens, at room temperature with Immersol W (Zeiss) between lens and coverslip. Using Zen software (Zeiss), live imaging of fluorescent proteins GFP and tdTomato was performed for 5 min with 5s interval on average. If GFP only, then the interval was 2s on average. Only movies obtained from larvae that were still alive and active after imaging were used for analysis. Each movie was then registered on ImageJ using StackReg plugin before analysis^72^.

### Quantification of Lifeact::GFP and Spir::GFP localization

New branches appearing in each movie were identified, and initiation sites were classified as having *de novo* actin polymer if a punctum of LifeAct::GFP intensity appeared at the site and grew in intensity until protrusion of the new branch. Initiation sites had “pre-formed” actin if motile blobs of LifeAct::GFP were observed to arrive prior to branch outgrowth (motile). Instances where a new branch grew from a site where a LifeAct::GFP punctum existed in the first frame of the movie were disregarded because the movie did not capture the origin of the actin punctum. Consistent with a previous report^16^, we observed only a single rare instance where a filopodium grew from an initiation site where LifeAct::GFP was not detected. Overall LifeAct::GFP expression in *spir* mutants appeared lower than in controls, consistent with our previous report for Spir knockdown with RNAi^23^, but it was certainly sufficient to monitor actin polymer at sites of branch initiation.

To measure the association of Spir::GFP labeled punctae to either a dendrite shaft, branch point, or tip, a custom routine was developed to extract this information from images acquired with a 20X objective (2 to 4 images stitched together where necessary using the Pairwise Stitching plugin^73^. From maximum intensity projections (MIP) of 10 c4da neurons (ddaC) labeled for the dendrite marker CD4-tdTom, signal from beyond the cell of interest was cleared manually, the background was subtracted, and a MIP mask created. This mask was then applied to the SpirFL::GFP channel. Pixels outside the mask were cleared. The MIP mask was skeletonized using Fiji’s skeletonization plugin^74^ to obtain the location of shafts, branch points, and tips. A script from the Neuroantomy update site was then used to determine which Spir::GFP punctae were associated with such skeleton features. When a Spir::GFP punctum was 3 µm or less (snap-to distance) from either a branch point or tip it was assigned to each, respectively. This snap-to distance was selected because from a representative Spir:GFP image, 80% of the punctae had a diameter of 3 µm or less. Additionally, no branch points or tips were 3 µm or less apart from one another. The number of punctae at branch points and tips was then divided by the total number of branch points and tips, respectively, to obtain the percent of branch points and tips that were occupied by Spir::GFP.

### Dendrite morphometry

For morphometry of c4da neurons (ddaC) at L3, images were acquired at 20X, with 1 image field usually sufficient to capture the entire dendritic field of a neuron. However, in some cases 2 images were stitched together using the Fiji plugin Pairwise Stitching to generate an image of the entire neuron^73^. Maximum intensity projections of confocal z-series image stacks were segmented as described previously^23^. Various Fiji plugins were used to determine branching characteristics of neurons; Sholl Analysis to generate Sholl profiles and extract Sholl related parameters^75^, Strahler Analysis to assess the number of terminal dendrites^75,76^, and Analyze Skeleton for total arbor length. Dendritic field was determined by fitting a polygon around the segmented neuron on Fiji, and measuring the area inside the shape^77^. For morphometry of ddaC neurons at L2 in time-lapse movies, the number of terminal dendrites/µm^2^ was calculated in the first frame of each movie. For the duration of each movie, the number of new branches/µm^2^/minute was calculated.

### Measuring dendrite dynamics

Pre-existing dendrites at the beginning of live imaging movies were analyzed for the dynamics of their terminal branches using MTrackJ^78^. Terminal branch length was compared by an observer every frame of each movie. Dendrites that extended, retracted, or did both, or remained static were recorded, and these numbers were expressed as the percentage of the total number of pre-existing dendrites at the beginning of each movie. Newly formed dendrites were tracked similarly, and their protrusive outgrowth was measured, where mean speed was the average of the speeds of growth for each frame (including extensions and pauses) until the branch stopped growing altogether, or retracted, or the movie ended. The proportion of time the branch paused during this period of saltatory outgrowth was calculated, as was the maximum length of its protrusion.

### Graphs and Statistics

Bar graphs and statistics were done using GraphPad Prism v9, and displayed data is expressed as mean ± standard error (SEM). Asterisks indicate the significance of P-values compared to the indicated control group (*P<0.05, **P<0.01, ***P<0.001, ****P<0.0001). For comparisons of more than two conditions, data were tested for normal distribution using D’Agostino & Pearson test and analyzed for statistical significance using one-way ANOVA (multiple groups) and Dunnett’s post-hoc tests were performed. For the repeated measures of western blots, Friedman test was applied then Dunn’s post-hoc tests. To compare two conditions, Student’s t-test (unpaired, two-tailed) was used.

## Supplemental Information

**Supplemental Figure S1:**
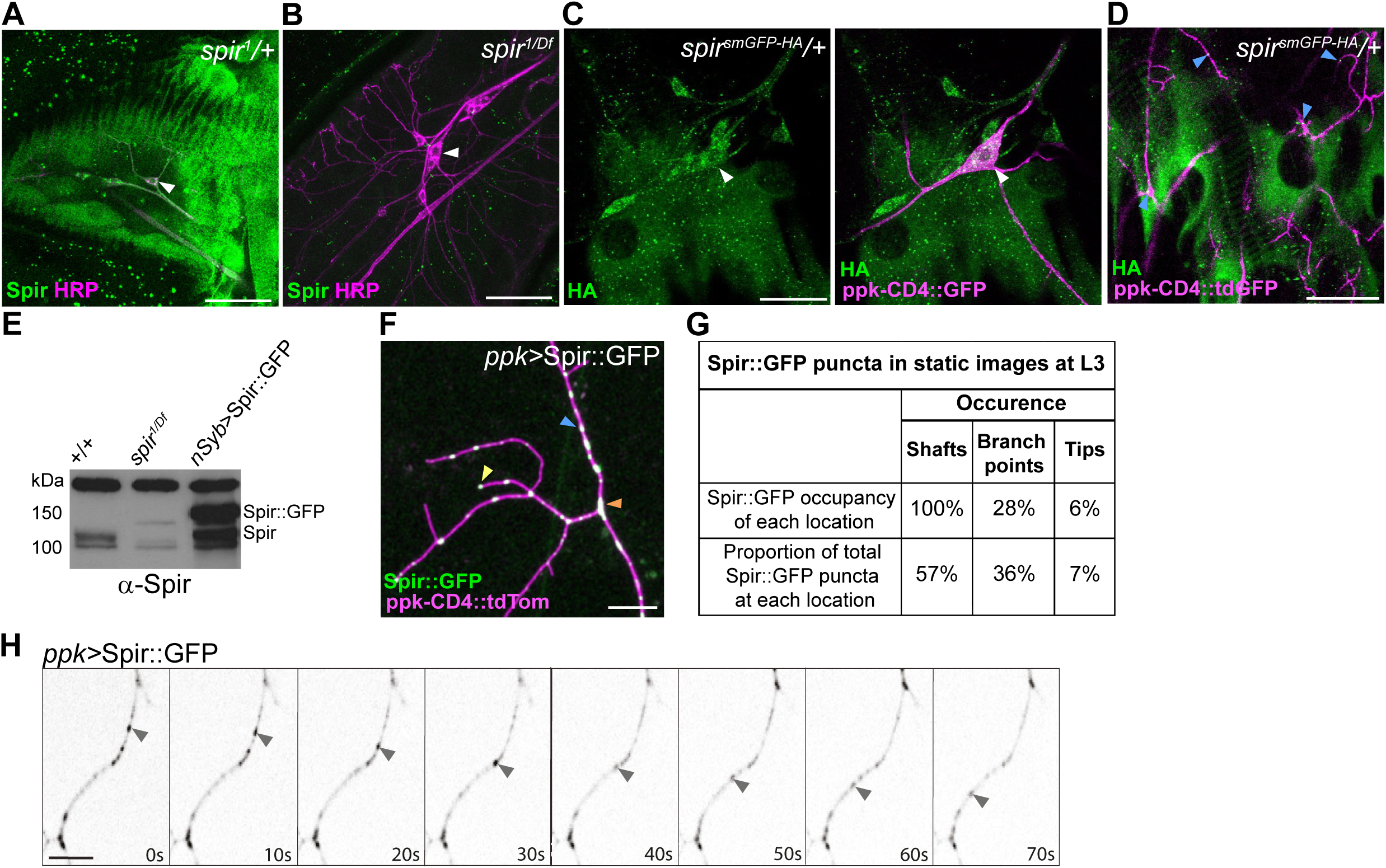
Spir expression and motility in dendrites (related to Figure 2). (A, B) IHC co-labeling for endogenous Spir and horseradish peroxidase (HRP) in the body wall of L3 larvae, showing Spir expression in a *spir^1^/+* heterozygote (A) but not in a *spir^1/Df^* hemizygous mutant (B). (C, D) IHC to detect HA-tagged Spir in a Spir-smGFP-HA larva, focusing on cell body of a c4da neuron (C), and distal dendrite arbors (D). (E) Anti-Spir western blot of lysates from dissected adult brains, comparing controls with *spir^1/Df^*mutants and with flies overexpressing Spir::GFP in all neurons with nSyb-Gal4. Expected mass of Spir::GFP is 142kDa, while that of Spir is 115kDa. The increased presence of Spir (115 kDa) upon overexpression of Spir::GFP likely arises from proteolytic cleavage of the GFP tag. (F) ppk- Gal4 driven Spir::GFP expression is detected with native GFP fluorescence in a c4da neuron labelled with tdTom in an L3 larva. In A-D and F, labelled c4da neuron cell bodies are marked (white arrowheads), as are puncta in dendrite shafts (blue arrowheads in D), branch points (orange arrowhead in F), and tips (yellow arrowhead F). Scale bars in A-D = 25um, F = 10 um. (G) Locations of Spir::GFP puncta observed in static images of c4da dendrite arbors at L3 (n=10 neurons). (H) Greyscale filmstrip (Supplemental Video 4) from a c4da neuron of an L2 larva, where the black arrowhead indicates progressive movement of a Spir::GFP-labelled punctum within a dendrite shaft. Scale bar: 5μm. Motile Spir::GFP puncta moved in both anterograde (64%) and retrograde (36%) directions from the cell soma, and sometimes were observed to switch from one to the other. Some puncta appeared to pass one another, while others merged. Motile Spir::GFP particles were found to travel 0.6 um/s on average, and achieved a maximal speed of 1.1 um/s.

**Supplemental Figure S2:**
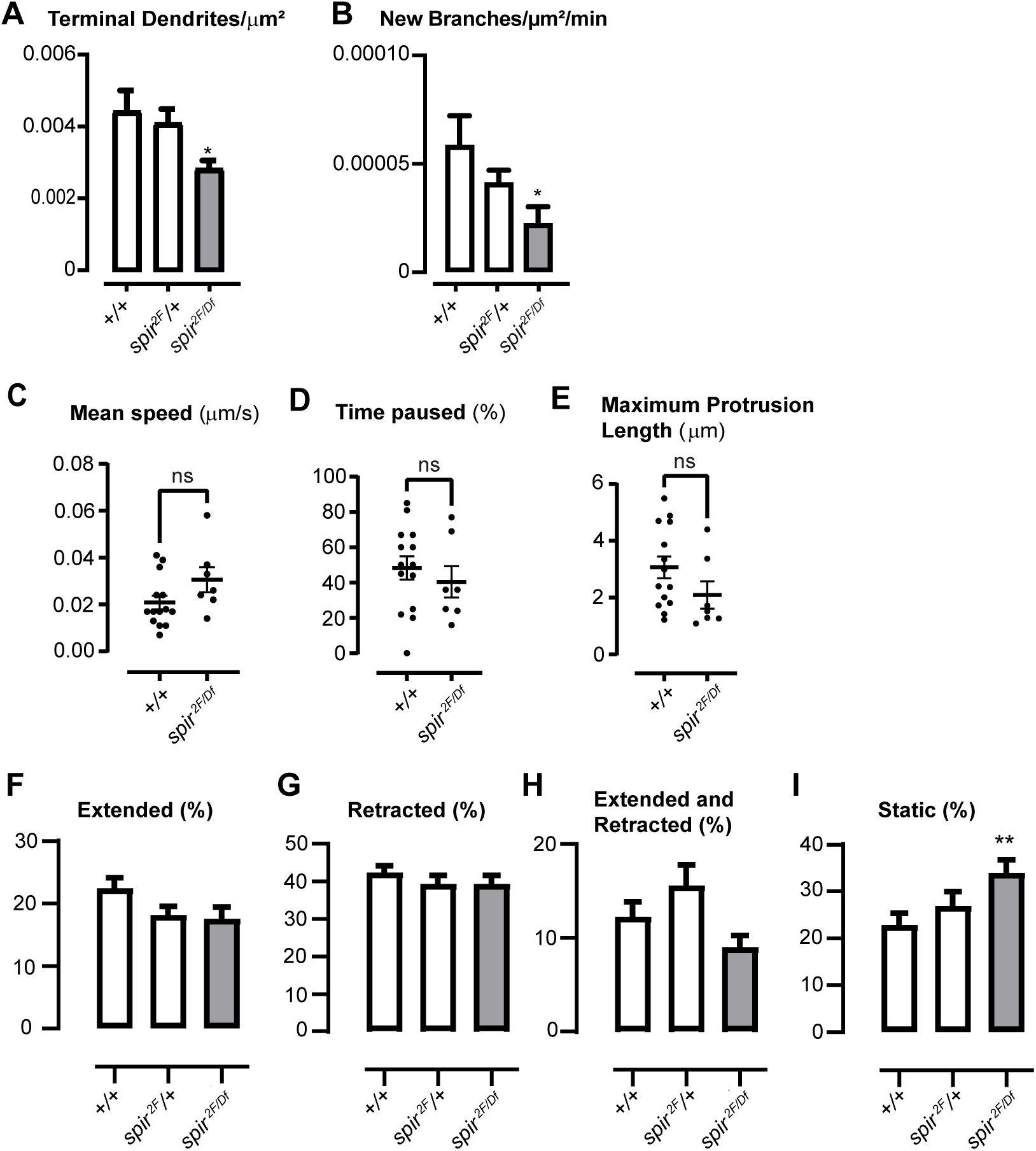
Quantification of dynamics of newly formed and pre-existing terminal branches (related to Figure 2). (A-B) Dendrite parameters from time-lapse movies of c4da neurons (labeled with ppkCD4::tdTom) in L2 larvae expressing LifeAct::GFP driven by *ppk-GAL4.* Graphs show mean ± SEM, comparing controls (+/+, n=11 movies) with *spir* heterozygotes (*spir^2F^*/+, n=8) and *spir* mutants (*spir^2F^*/*spir^Df^*, n=11). Asterisks indicate significant changes compared to +*/+* controls. (A) Number of terminal branches per µm^2^ of area in first frame of each movie (ANOVA (F(2, 27) = 4.079, p = 0.0283). (B) Number of new branches/µm^2^/min in each movie (ANOVA (F(2, 27) = 3.265, p = 0.0537). (C-E) Protrusive growth characteristics of nascent branches in controls (+/+, n=14 movies) and *spir* mutants (*spir^2F^*/*spir^Df^*, n=7) in L2 larvae expressing LifeAct::GFP driven by *ppk-GAL4*, including mean speed (C, t-test (t(21) = 1.757, p = 0.0951), time paused (D, t-test(t(21) = 0,7025, p = 0.4909), and maximum protrusion length (E, t-test (t(21) = 1.516, p = 0.1459). Graphs show scatter plot for each branch, with mean ± SEM indicated (ns, not significant). (F-I) For branches that were already present in the first frame of each movie, quantification of percentage of pre-existing branches that extended (F, ANOVA (F (2, 62) = 2.482, p=0.0919), retracted (G, ANOVA (F (2, 62) = 0.6473, p=0.5269), extended and retracted in the same movie (H, ANOVA (F (2, 62) = 4.058, p=0.0221) or remained static (I, ANOVA (F (2, 62) = 4.389, p=0.0165). Results in F-I are combined for neurons that expressed LifeAct-GFP and those that did not, since LifeAct-GFP does not affect the number or dynamics of pre-existing branches. Graphs show mean ± SEM, comparing controls (+/+, n=21 movies) with *spir* heterozygotes (*spir^2F^*/+, n=18) and *spir* mutants (*spir^2F^*/*spir^Df^*, n=26).

**Supplemental Figure S3:**
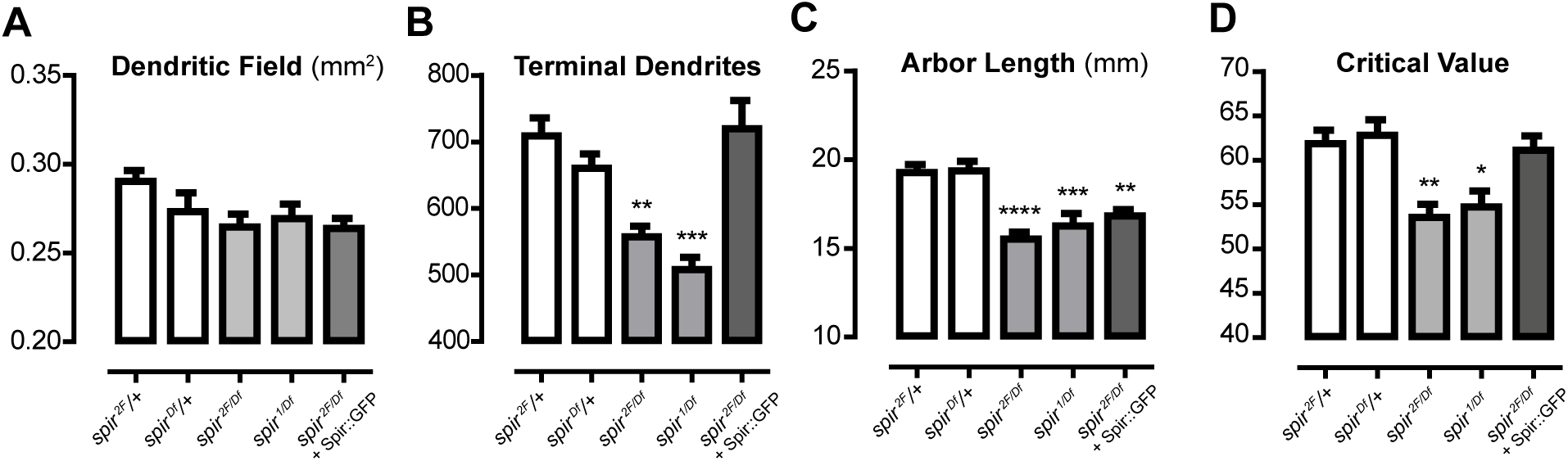
Dendrite parameters for c4da neurons in L3 larvae (related to Figure 3). (A-D) This is the same dataset shown in Fig. 3, except it was adjusted to control for dendritic field size by removing from the analysis four *spir^2F/Df^* c4da neurons with unusually small arbors (n=11). Asterisks indicate significant changes compared to *spir^2F^*/+ heterozygous controls. (A) Dendritic field per neuron (mm^2^) (ANOVA (F(4, 66) = 1.912, p = 0.1189. (B) Total number of terminal dendrites per neuron (ANOVA (F(4, 66) = 9.328, p < 0.0001). (C) Total length of dendrite arbor per neuron (mm) (ANOVA (F(4,66) = 12.03, p < 0.0001). (D) Sholl critical value (ANOVA (F(4, 66) = 6.914, p = 0.0001).

**Supplemental Figure S4:**
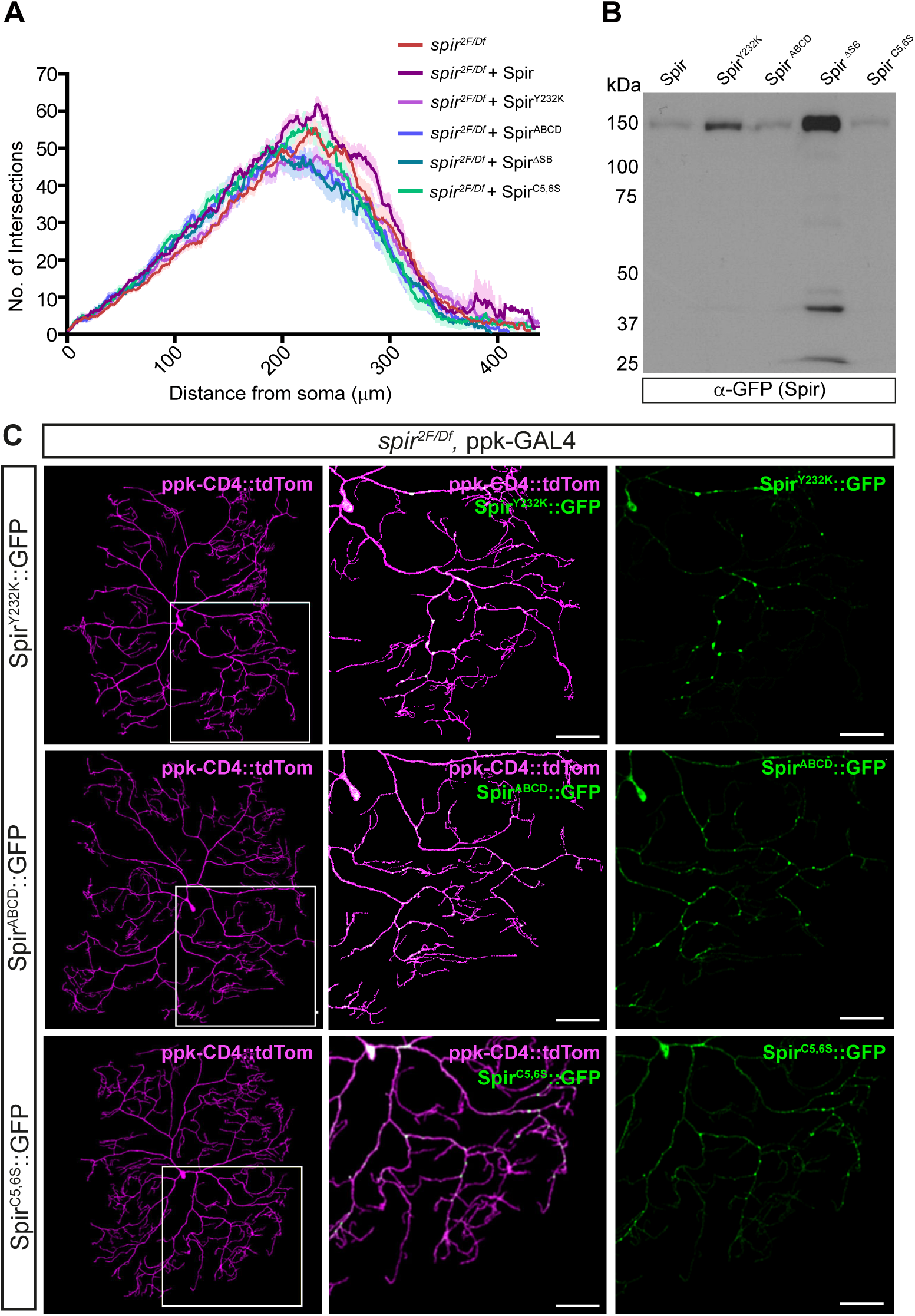
Function, expression, and distribution of Spir::GFP bearing domain-specific mutations (related to Figure 4). (A) Sholl profiles for entire c4da neurons labelled with ppk-CD4::tdTom. (B) Anti-GFP western blot from adult heads of animals overexpressing in all neurons either intact Spir::GFP or Spir::GFP bearing domain-specific mutations. One of three replicate blots, this blot is distinct from the one shown in Fig. 4C and is shown in its entirety. The expected molecular weight of Spir::GFP is 142 kDa, and that of GFP is 27 kDa, and so proteolytic cleavage of GFP is not apparent for most constructs. (C) *ppk-GAL4* drives the expression of Spir^Y232K^::GFP or Spir^ABCD^::GFP or Spir^C5,6S^::GFP in a *spir* mutant background. Punctate distribution of Spir::GFP is maintained for each of these mutations. White boxes in left panels indicate areas magnified in panels to their right. Scale bar=50μm.

**Supplemental Figure S5:**
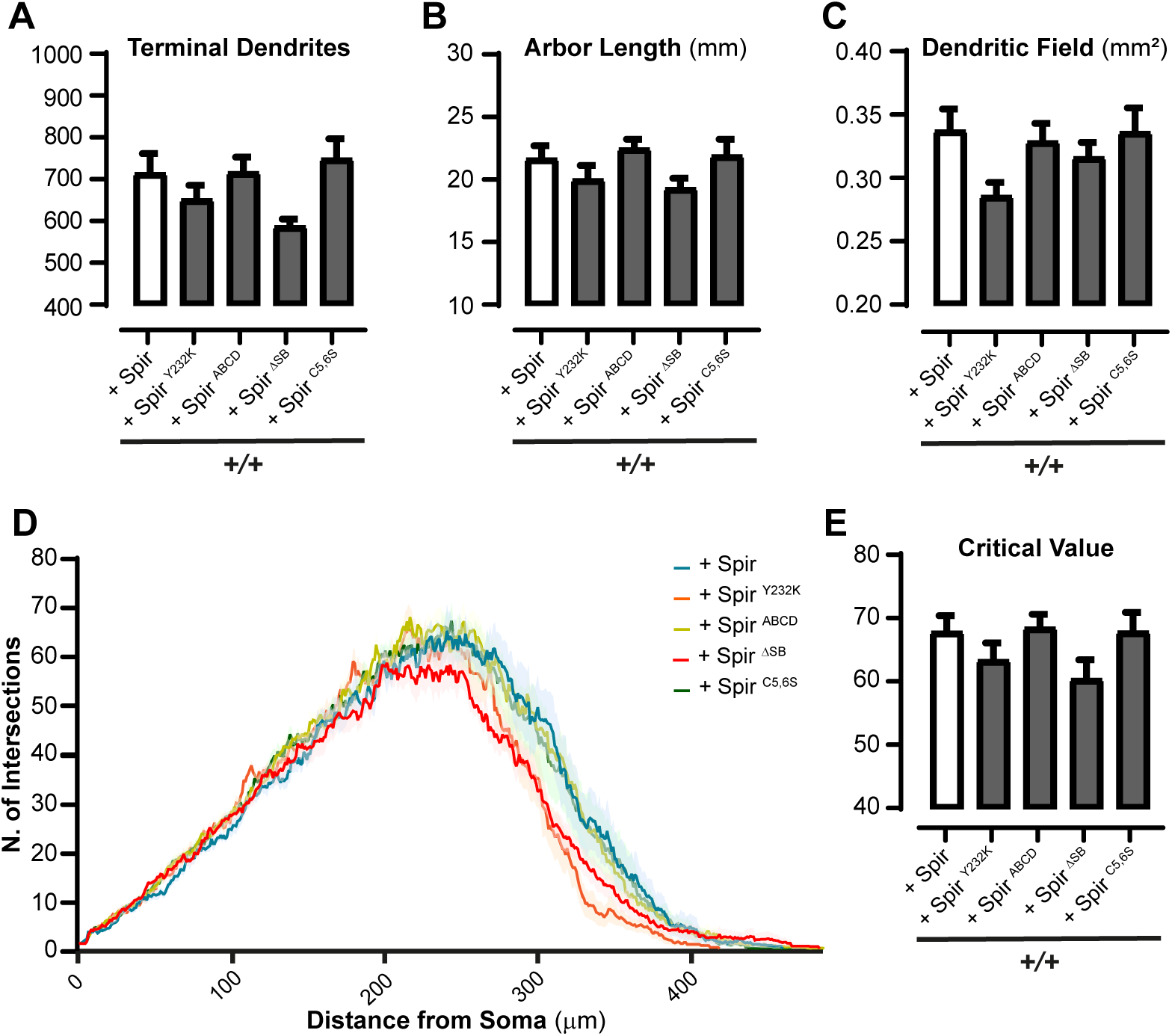
Effects of overexpression of Spir domain-specific mutations on c4da dendrite arborization in wild-type (+/+) animals (related to Fig. 4). (A-E) Dendrite parameters (mean ± SEM) in wild-type L3 larvae (+/+) expressing Spir::GFP (“+ Spir”), or Spir:GFP with domain-specific mutations (n=8-10 for each genotype). Asterisks indicate significant changes compared to + Spir controls (white bar). All animals in this study bore ppk- Gal4 and ppk-CD4::tdTom transgenes. (A) Total number of terminal dendrites per neuron (ANOVA F(4, 40) = 3.141, p = 0.0245). (B) Total length of dendrite arbor per neuron (mm) (ANOVA F(4, 40) = 1.783, p = 0.1513). (C) Dendritic field per neuron (mm^2^) (ANOVA F(4, 40) = 1.843, p = 0.1397). (D) Sholl profiles. (E) Sholl critical value (ANOVA F(4, 40) = 1.716, p = 0.1655).

**Supplemental Figure S6:**
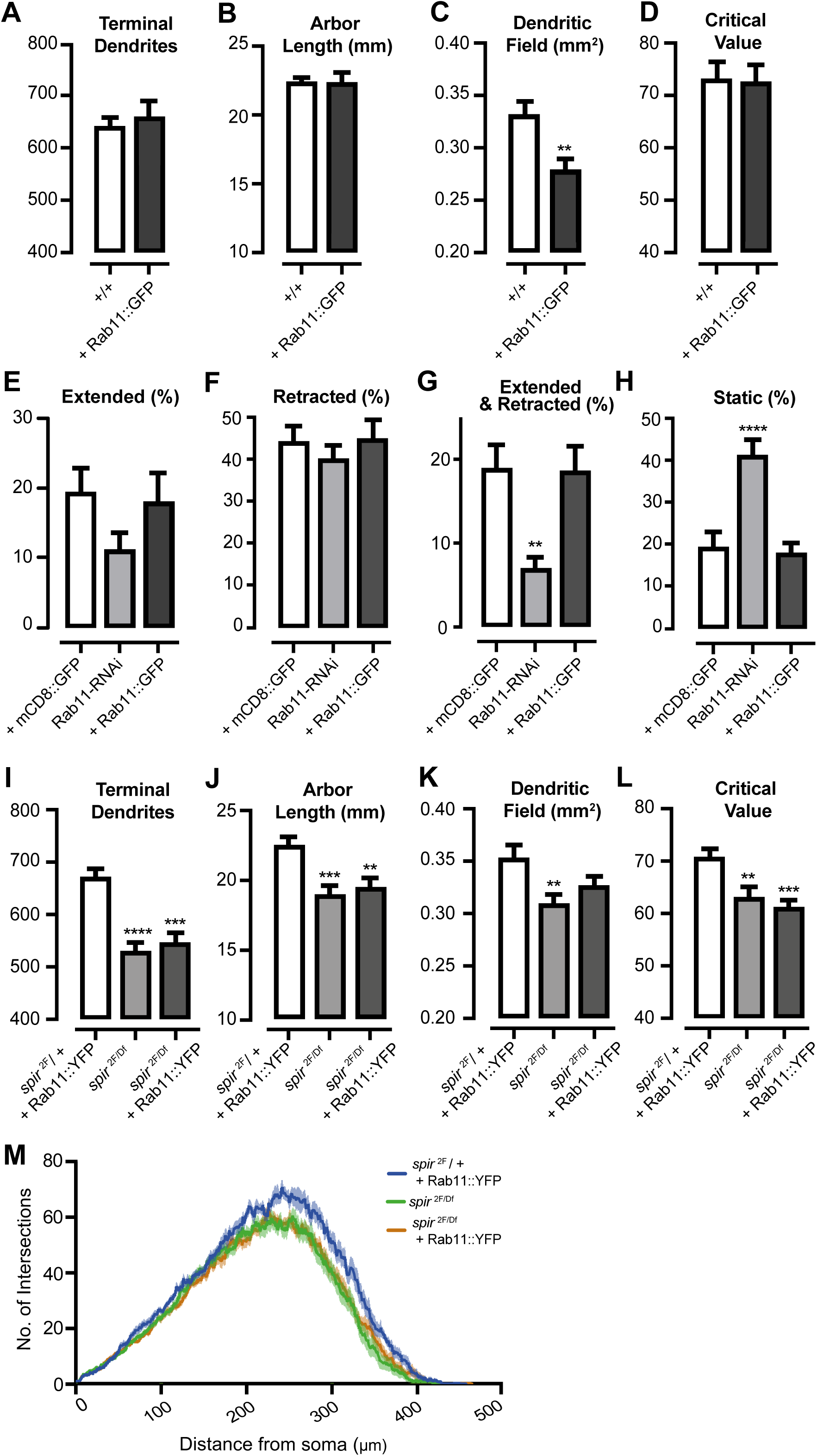
Effects of Rab11 on dendrite dynamics at L2 and dendrite arborization at L3 (related to Figure 5). (A-D) Dendrite parameters (mean ± SEM) in control L3 larvae (+/+) and ppk-GAL4-driven Rab11::GFP (n=10 for each genotype). Asterisks indicate significant changes compared to controls. (A) Total number of terminal dendrites per neuron (t-test, t(18) = 0.5270, p = 0.6046). (B) Total length of dendrite arbor per neuron (mm) (t-test, t(18) = 0.0571, p = 0.9550). (C) Dendritic field per neuron (mm^2^) (t-test, t(18) = 3,551, p = 0.0023). (D) Sholl critical value (t-test, t(18) = 0.1905, p = 0.8510). (E-H) Dendrite parameters from time-lapse movies of c4da neurons (labeled with ppkCD4::tdTom) in L2 larvae. Graphs show mean ± SEM, comparing controls expressing mCD8::GFP (n=10 movies) with Rab11-RNAi driven by *ppk-GAL4* (n=11) and Rab11::GFP driven by *ppk-GAL4* (n=10). Asterisks indicate significant changes compared to controls. For branches that were already present in the first frame of each movie, quantification of percentage of pre-existing branches that extended (E, ANOVA (F (2, 28) = 2.301, p = 0.1188), retracted (F, ANOVA (F (2, 28) = 0.5551, p = 0.5802), extended and retracted in the same movie (G, ANOVA (F (2, 28) = 8.343, p = 0.0014) or remained static (H, ANOVA (F (2, 28) = 16.50, p<0.0001). (I-M) L3 dendrite parameters (mean ± SEM) in control *spir*^2F/+^ heterozygotes expressing Rab11::YFP, *spir^2F/Df^* mutants with or without Rab11::YFP. (n=10 for each genotype). Asterisks indicate significant changes compared to controls (*spir*^2F/+^ heterozygotes + Rab11::YFP). (I) Total number of terminal dendrites per neuron (ANOVA (F(2, 27) = 16.21, p < 0.0001). (J) Total length of dendrite arbor per neuron (mm) (ANOVA F(2, 27) = 12,10, p = 0.0002). (K) Dendritic field per neuron (mm^2^) (ANOVA (F(2, 27) = 4.941, p = 0.0148). (L) Sholl critical value (ANOVA (F(2, 27) = 8.764, p = 0.0012). (M) Sholl profiles.

**Supplemental Figure S7:**
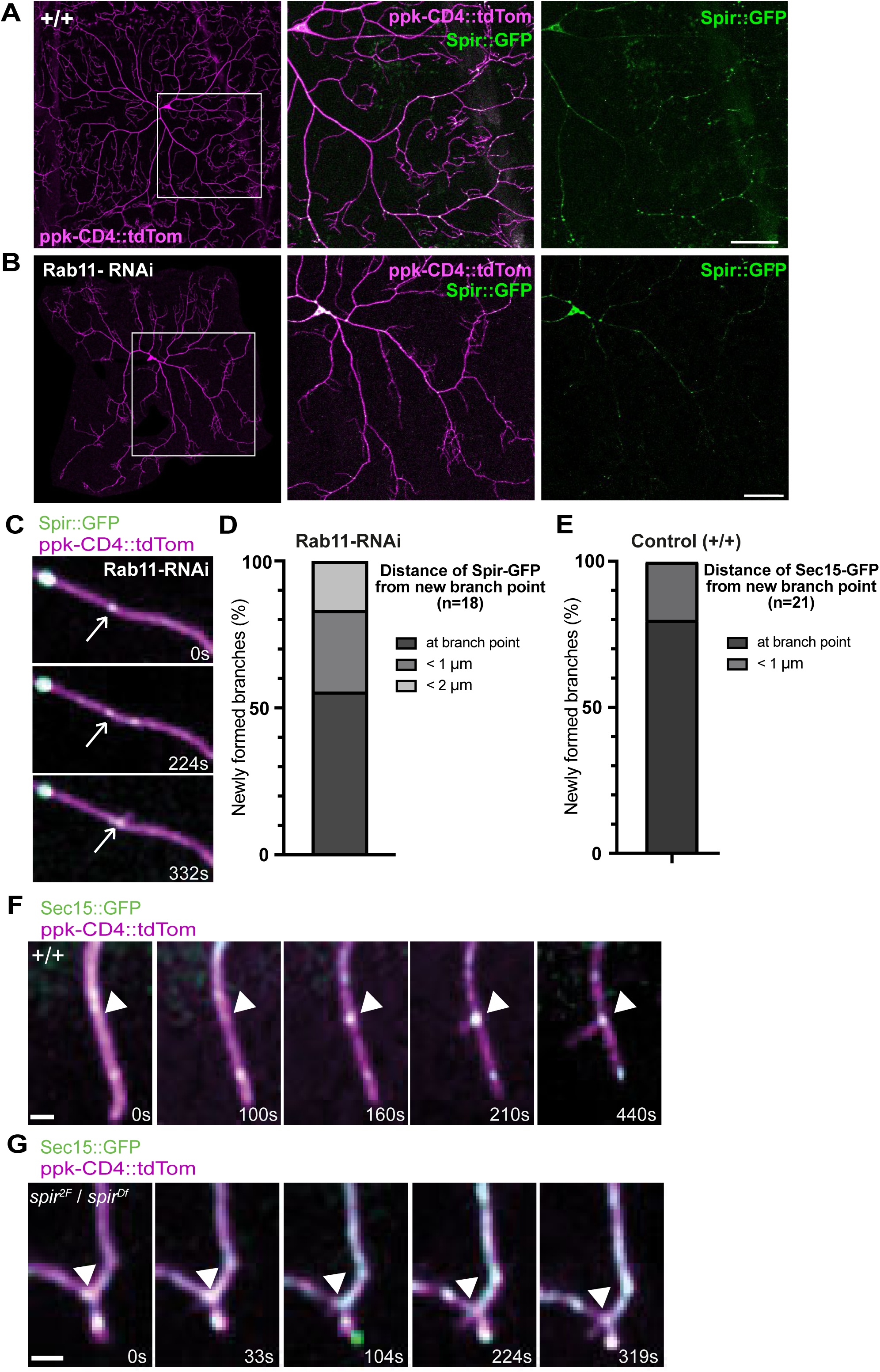
Expression, and distribution of Spir::GFP and Sec15::GFP in c4da neurons (related to Figures 5 and 6). (A, B) Distribution of Spir::GFP in c4da neurons of L3 larvae in controls (+/+, A) and upon Rab11 RNAi knockdown in c4da neurons with *ppk- Gal4* (B). White boxes in left panels indicate areas magnified in panels to their right. Upon knockdown of Rab11 in c4da neurons, Spir::GFP retained punctate distribution within dendrite arbors. Scale bars=50μm. (C) Filmstrip (Supplemental Video 8) of Rab11 knockdown L2 larva, where Spir::GFP (arrows) was observed at the initiation site of a new dendrite branch. (D) In Rab11 knockdown c4da neurons, the distance of the closest Spir::GFP puncta from the site of branch initiation (n=18). (E) In control (+/+) c4da neurons, the distance of the closest Sec15::GFP puncta from the site of branch initiation (n=21). (F) Filmstrip (Supplemental Video 9) of control (+/+) L2 larva, where Sec15::GFP (arrowhead) pre-exists at the initiation site of a new dendrite branch. (G) Filmstrip (Supplemental Video 10) of *spir^2F/Df^* mutant L2 larva, with Sec15::GFP (arrowhead) at a rare branch initiation site.

**Supplemental Video 1 – corresponding to Fig. 1C**. 1 frame/7.2s.

**Supplemental Video 2 - corresponding to Fig. 1D**. 1 frame/4.4s.

**Supplemental Video 3 - corresponding to Fig. 1E**. 1 frame/6.5s.

**Supplemental Video 4 - corresponding to Supplemental Fig. S1H**. 1 frame/∼2s.

**Supplemental Video 5 - corresponding to Fig. 2A**. 1 frame/∼4s.

**Supplemental Video 6 - corresponding to Fig. 2B**. 1 frame/∼7s.

**Supplemental Video 7 - corresponding to Fig. 5G** 1 frame/∼6s.

**Supplemental Video 8 - corresponding to Supplemental Fig. S7C**. 1 frame/∼7s.

**Supplemental Video 9 - corresponding to Supplemental Fig. S7F**. 1 frame/∼7s.

**Supplemental Video 10 - corresponding to Supplemental Fig. S7G**. 1 frame/∼7s.

